# Terminal conjugation enables nanopore sequencing of peptides

**DOI:** 10.1101/2025.11.12.687956

**Authors:** Justas Ritmejeris, Xiuqi Chen, Bauke Albada, Cees Dekker

## Abstract

Nanopore sequencing of peptides holds great promise for single-molecule proteomics, but robust conjugation strategies to adapt native peptides for motor-enzyme–driven translocation have yet to be developed. Here, we establish terminally directed DNA–peptide conjugation chemistry strategies that expand the applicability of nanopore sequencing beyond synthetic model systems to natural peptides. At the N terminus, omniligase catalyzes rapid and peptide ligation of a DNA handle under mild conditions. At the C terminus, photoredox decarboxylative ligation introduces a bioorthogonal linker that enables CuAAC-mediated DNA attachment that ensures proper stretching and translocation of short peptides through the nanopore. Our study reveals that long peptides can be sequenced with single-end conjugation, while short or neutral peptides require threading tails. Positively charged peptides cannot be translocated under the same electric field but can be sequenced after charge neutralization. The data demonstrate controlled nanopore readouts of peptides that differ widely in length, charge, and sequence. This framework establishes a versatile chemical foundation for adapting natural peptides to nanopore sequencing, advancing single-molecule proteomic analysis.

## Introduction

Nanopore sequencing has transformed genomics and transcriptomics by providing single-molecule analysis of DNA and RNA^1–3^. Extending this capability to peptides and proteins would enable direct observation of proteoforms, post-translational modifications, and molecular heterogeneity – all at a single-molecule level inaccessible to mass spectrometry^4,5^. However, protein sequencing faces various significant challenges. For example, peptides lack the uniform negative charge and predictable structure of nucleic acids, preventing reliable electrophoretic capture and controlled translocation through nanopores. Furthermore, free translocation of proteins yields very short residence times in the nanopore^6^, which prevents their accurate analysis. Motor enzymes have proven essential for achieving nanopore sequencing by enforcing directional movement and slowing analyte translocation to the (sub)second timescales^7,8^. The protein unfoldase ClpX for example was shown to induce translocation of full-length proteins^9–11^. However, ClpX requires specific recognition tag for substrate engagement and often displays irregular multi-residue step sizes^12^, which limits its general applicability for high-resolution sequencing.

DNA–peptide conjugation has emerged as an alternative strategy that incorporates the exceptional processivity and precision of DNA motor enzymes to control analyte movement through nanopores^13–15^. By attaching DNA handles to peptides, molecular motor proteins like DNA helicases can control peptide translocation while maintaining their native stepping behavior on the DNA strand (Fig.1A). For example, the helicase Hel308 provide a uniform ∼0.3 nm step size along DNA with minimal backtracking, enabling 2 steps per base resolution in nanopore DNA sequencing platforms^16^. Proof-of-concept studies demonstrated single–amino acid resolution in peptide translocation^13^ as well as PTM detection^17,18^ using this approach. However, these studies relied exclusively on synthetic peptides with pre-installed cysteines or bioorthogonal handles for conjugation to DNA^13–15,17–19^, limiting its general applicability. Such requirements preclude the analysis of native peptides derived from biological samples, where modification sites cannot be predetermined.

**Figure 1.**
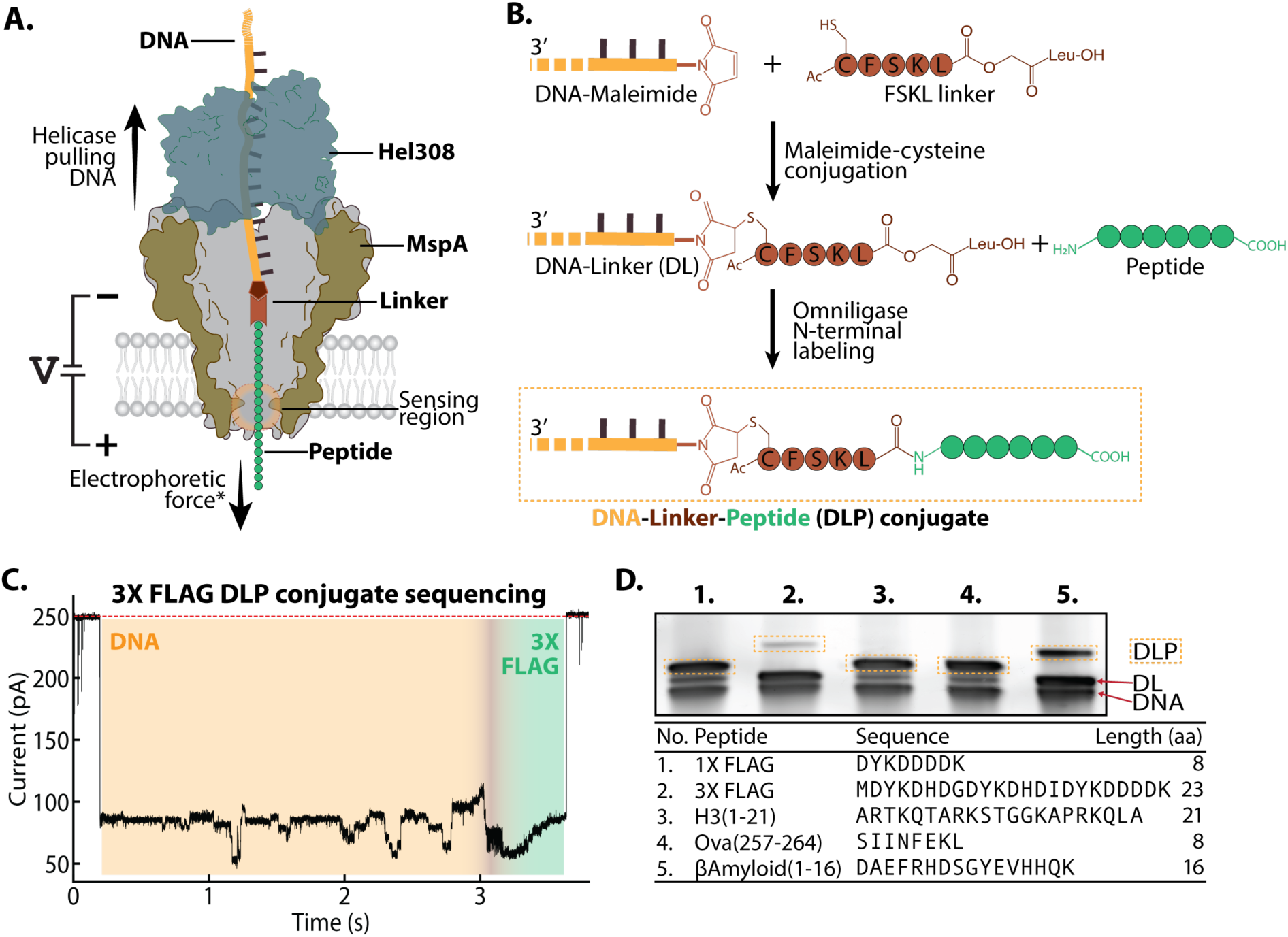
Omniligase enables site-specific N-terminal DNA conjugation of peptides compatible with nanopore sequencing. (A) Schematic depiction of nanopore sequencing of DLP constructs. Under an applied voltage (+180 mV), negatively charged DLP molecules are electrophoretically driven into MspA-M2(N93D) pores until stopped by the Hel308 motor enzyme, which subsequently pulls the DNA back through the pore. Corresponding ionic current changes can be assigned to the DNA, linker, and peptide segments as they traverse the sensing region. (B) Schematic depiction of the N-terminal conjugation strategy. First, a DNA oligonucleotide with a maleimide group is coupled to a cysteine-containing FSKL linker. The resulting DNA–linker (DL) conjugate is ligated to the peptide of interest using omniligase enzyme, yielding a DNA–linker–peptide (DLP) construct. (C) Representative nanopore trace of a urea-PAGE-purified 3X FLAG DLP construct recorded using MinION flowcells containing MspA-M2(N93D) pores, showing a DNA region followed by a gradual transition into the linker and then into the peptide segment. Sequencing was performed in cis buffer containing 500 mM KCl, 10 mM HEPES, 10 mM MgCl₂, and 1 mM ATP. (D) Urea-PAGE (10%) analysis of DLP formation with various peptides. Distinct band shifts confirmed peptide attachment to the DL with varying ligation efficiencies.

Several chemical and enzymatic methods have been explored for terminal peptide modifications, but each of these is subject to specific sequence requirements or reaction conditions. At the N terminus of peptides, a wide variety of strategies are available^20^. Native chemical ligation and transamination-based methods are highly specific in controlled contexts but their applicability is restricted by sequence dependence, i.e. it requires specific amino acids at the N terminus^21–23^. N-hydroxysuccinimide (NHS)-esters are widely used but typically modify not merely the N-terminal but also all lysine side chains^24^. 2-pyridinecarboxaldehyde (2-PCA) reagents provide a versatile approach that disfavors only proline in the second residue, but it requires extended reaction times and a high excess of reagents^25–28^. At the C terminus, enzymatic strategies using Sortase A enable efficient ligation but this approach relies on the engineered LPXTG motif, limiting its generalization^29,30^. More broadly, bifunctional cross-linking chemistries can connect DNA to peptides but often yield heterogeneous mixtures^31,32^. Despite progress in protein labeling strategies, no existing method provides a universal, and terminal-specific strategy for DNA attachment at both termini of natural peptides.

Here we developed terminal conjugation strategies of adapting native peptides for nanopore sequencing. We integrate enzymatic N-terminal ligation with chemical C-terminal functionalization to create DNA–peptide–DNA conjugates. Enzymatic N-terminal ligation was achieved with omniligase, which can couple peptides to a DNA-linker intermediate within minutes under mild conditions^33^. Although broadly sequence-tolerant, this reaction is influenced by the two residues at the N-terminal of the target peptide^34^, and hence we developed a strategy to avoid this dependence. At the C terminus, photoredox-mediated decarboxylation was used to selectively activate the terminal carboxylate, enabling installation of an alkyne-containing linker under visible-light irradiation^35,36^. Subsequent copper-catalyzed azide-alkyne cycloaddition (CuAAC) coupling^37^ of an azide-functionalized DNA tail provided constructs with enhanced capture and stretching properties. Together, these chemistries offer orthogonal and broad-sequence compatibility to both termini of natural peptides.

By combining these chemical strategies, we analyze diverse peptides ranging in length from 8–26 residues with highly varying charge profiles. We establish key design rules for nanopore sequencing approaches. Specifically, (i) longer peptides (>∼ 20 amino acids) with a net negative charge can be analyzed using N-terminal conjugation alone, (ii) shorter peptides require an additional C-terminal DNA for stable capture and translocation, and (iii) chemical neutralization of positively charged peptides enables sequencing of cationic peptides that are otherwise excluded by electrostatic repulsion. These terminal-conjugation strategies provide the first general method for adapting native peptides to motor-driven nanopore sequencing, establishing a chemical foundation for single-molecule proteomic applications.

## Results and Discussion

### Omniligase enables rapid N-terminal DNA conjugation across diverse peptide sequences

To systematically evaluate peptide conjugation and sequencing performance, we assembled a panel of 11 peptides spanning 8–26 residues (Supplementary Table 1). This captures a representative range of peptides of various length, charge, and sequence contexts encountered in natural samples.

To enable DNA motor-driven nanopore sequencing of natural peptides, a DNA handle was introduced at the N-terminus of the peptide. Omniligase catalyzes scarless peptide bond formation between an N-terminal amino group of a substrate peptide and a small acyl donor peptide carrying a C-terminal –OCam-LeuOH activating ester^34^. To adapt this chemistry for DNA–peptide conjugation, we synthesized a DNA oligonucleotide bearing a 5′-maleimide group and coupled it to a pentapeptide Ac-CFSKL-OCam-LeuOH (linker peptide) via maleimide–cysteine chemistry (Figure 1B and Supplementary Figure 1). The resulting DNA–linker (DL) (Supplementary Figure 2) served as the acyl donor for omniligase-mediated ligation with peptides of interest, yielding DNA–linker–peptide (DLP) conjugates (Figure 1B, bottom). Such DLPs can then be subject to sequencing with the MspA nanopore/Hel308 helicase system. Once the DLP is bound by the Hel308 in bulk, it is electrophoretically driven into the pore and halted as the helicase is drawn on top of the MspA nanopore (Figure 1A). The partial blockade of the nanopore constriction region (i.e. its narrowest part) produces step-like ionic current signatures as the helicase pulls the molecule upwards against the electric field (Figure 1C). The peptide signals immediately follow the well-characterized DNA signals, allowing for sensitive characterizing of the peptide properties^13^.

We performed omniligase-ligation-based preparation of DLPs and evaluated the ligation efficiency by urea PAGE. Distinct band shifts confirmed successful conjugation (Figure 1D, Supplementary Figure 3). As the reaction reached maximal yield within 5 minutes at room temperature (see H3(1-21) peptide data in Supplementary Figure 4), this method appears to be compatible with time- or temperature-sensitive samples^38^. Conjugation yields depended on peptide sequence and varied from 23% for the 3X FLAG peptide to 67% for the H3(1-21) peptide, with averages determined from three independent reactions. Consistent with prior reports, efficiency depended on the N-terminal residues of the acceptor peptide^34,39,40^. Additional factors such as charge profile, length, and peptide solubility however also influence ligation efficiency. To validate that the shifted bands corresponded to ligated peptides, 1X and 3X FLAG DLPs were isolated by affinity purification using anti-FLAG resin, yielding a clear conjugate band at the same location (Supplementary Figure 5). All DLP conjugates were purified by cutting the bands from the acrylamide gel before nanopore sequencing. An example is given in Figure 1C, which shows that the 3X FLAG peptide produced a distinct signal pattern after the DNA. At the boundary between DNA and peptide, we observed a period of rapid signal fluctuation, which likely relates to the movement of the opposing charges on the peptide (positively charged lysine and negatively charged aspartate) across the pore. Further into the peptide segment, the signals exhibited a reproducible signature with a slightly rising feature before returning to open pore level, when the DLP was ejected from the nanopore. With wide compatibility of peptide sequences for the omniligase^34^, this workflow creates a facile conjugation reaction for labeling N-terminus of various peptides.

### Single-terminal conjugation allows for robust measurement of long peptides but not short ones

Next, we examined the range of peptide lengths possible for omniligase-facilitated nanopore sequencing. Encouragingly, a large fraction of long peptide sequencing events (23–26 residues, Figure 2A) showed reproducible ionic current patterns with each variant displaying a characteristic signal profile that enabled discrimination (Figure 2B, Supplementary Figure 6). The maximal peptide read length in MspA corresponds to the distance between the helicase docking site and the sensing region, which is about 10 nm^13,41^. For DLPs, this distance covers the FSKL-linker (3.3 nm) and the first ∼17 amino acids (6.7 nm) if the molecule would adopt a linear stretched configuration. Accordingly, C(EGSG)₅EF and C(EGSG)₆Y, which share the first 22 residues, produced nearly identical signals (Figure 2B). However, note that the peptide does not adopt a straightened conformation due to insufficient stretching with low density of negative charge (Figure 2C, left), and hence the influence of the residues can be more distant than for a fully stretched linear peptide^13,42^. This also likely contributed to some signal deviations in the region with the same peptide sequence.

**Figure 2.**
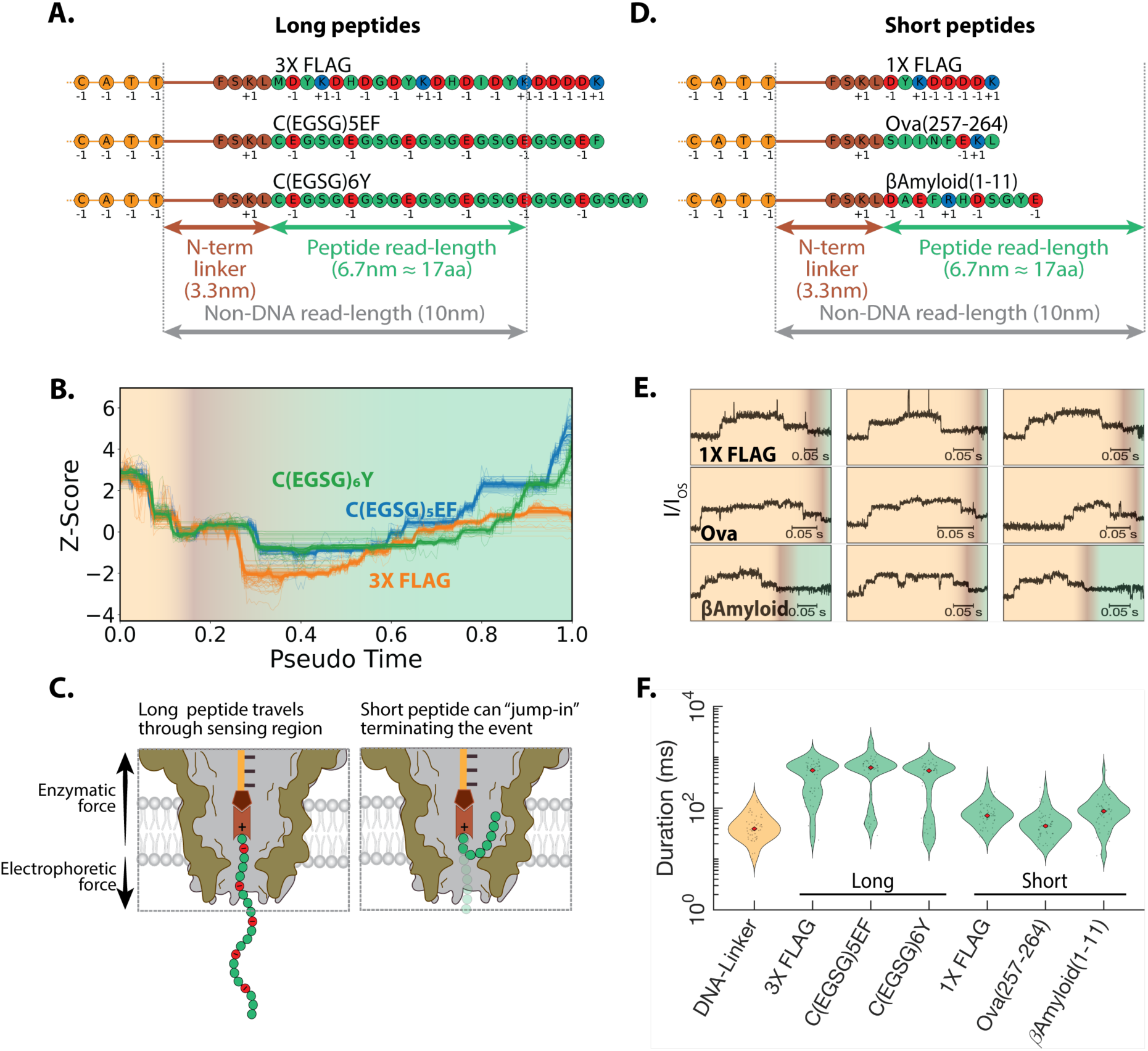
Nanopore sequencing of DLP conjugates reveals robust readout for long peptides but inconsistent signals for short peptides. (A) Schematic of DLP constructs containing long peptides of interest (23–26 amino acids) that are to be read in nanopore sequencing. The conjugates consist of DNA (orange), an FSKL linker (brown), and peptides with neutral (green), with negatively (red) or positively (blue) charged residues. Nanopore read length is primarily determined by the distance between the Hel308 docking site and the sensing region (10 nm for MspA). (B) Dynamic time warping (DTW) alignment of peptide-region signals for long-peptide DLP constructs: 3X FLAG (orange, N = 110), C(EGSG)₅EF (blue, N = 97), and C(EGSG)₆Y (green, N = 63). Gradient colors indicate transition from DNA (light yellow) to linker (brown) and peptide (green) regions. (C) Schematic depiction of a DNA motor enzyme pulling a negatively charged DLP conjugate through MspA. Electrophoretic force drives the construct into the pore, while the stronger enzymatic force ratchets it upward, resulting in a stretched conjugate that traverses the sensing region in a controlled fashion, while the bottom section behaves as an anchor to prevent premature event termination (left); for short-peptide sequencing, the positively charged linker and heterogeneous peptide sequence can enter the nanopore vestibule during sequencing event, prematurely terminating events and generating inconsistent signals at the peptide segment (right). (D) Schematic of DLP constructs containing short peptides of interest (8–11 amino acids). (E) Representative nanopore sequencing timetraces of short-peptide DLP constructs showing the transition from DNA (light yellow) to linker (light brown) to peptide (light green) regions. Scale bars represent individual event durations in seconds. Ion currents (I) are normalized to the local unblocked open-state pore current (I_OS_). All plots display the 0.3 – 0.65 I/I_OS_ range. (F) Duration of the peptide segment in the nanopore constriction, for different urea-PAGE-purified DLP constructs. DNA–linker (yellow) serves as a control to identify events lacking a peptide signal (median = 43 ms). Long-peptide constructs (3X FLAG, C(EGSG)₅EF, and C(EGSG)₆Y) show extended peptide-region dwell times (medians: 575, 635, and 562 ms, respectively). Short-peptide constructs (1X FLAG, Ova(257–264), and βAmyloid(1–11)) show markedly shorter durations (72, 44, and 89 ms, respectively). n=65 events per construct were used for analysis.

By contrast, short peptides (8–11 residues) (Figure 2D) were harder to detect and did not yield reproducible ionic current patterns, with individual events showing inconsistent signals (Figure 2E, Supplementary Figure 7). Although their length could in principle lead to a full translocation through the sensing region before helicase disengagement, most peptide variants displayed only very short signals, reducing a likelihood of precise peptide identification. Instead of consistent, stepwise signal traces, short peptide constructs frequently terminated prematurely and rarely demonstrated robust peptide signatures after the DNA trace, suggesting an unstable movement of the peptide across the nanopore constriction. We attribute this variability to the short length of the peptide within the pore (Figure 2C, right), which causes instability during translocation, and makes the peptide sensitive to thermal fluctuations as well as to interactions with the nanopore surface. This effect may be further amplified by a positively charged lysine residue in the linker, which reduces the effective electrophoretic stretching force acting on the peptide. In this regime, the electrophoretic force is insufficient to maintain tension across the molecule. This instability is also reflected in the peptide-region signal durations (Figure 2F), which showed markedly shorter dwell times (44-89 ms) for short-peptide constructs – closely matching the dwell time of DNA-linker events (43 ms) – in contrast to the longer peptides (562-635 ms). These results indicate that short peptides occupy the sensing region only briefly before escaping the nanopore.

To exclude purification artifacts as the source of variability, we compared urea-PAGE-purified samples (which often showed less pronounced DLP separation from DL for shorter peptides) with affinity-purified 1X and 3X FLAG DLPs obtained using anti-FLAG resin (Supplementary Figure 5). In both cases 1X FLAG peptide generated short signals while 3X FLAG peptide produced reproducible longer events with shorter events occurring only occasionally (Supplementary Figure 8).

These findings suggest that single-end conjugation requires a long peptide with net negative charge for consistency in nanopore readings. Shorter peptides fail to translocate orderly within the pore and yield events that often contain only the DNA segment, while the peptide portion is generally absent or indistinct (Supplementary Figure 7). This motivated the attachment of C-terminal threading tails, which should stabilize short peptides in the nanopore and extend the range of sequences compatible with nanopore sequencing, as discussed next.

### Conjugation at both peptide termini allows nanopore sequencing of a broader range of peptides

To extend the applicability of DNA–peptide conjugates to short peptides, we developed a C-terminal modification strategy that installs a negatively charged threading tail, yielding DNA-linker-peptide-linker-DNA (DLPLD) conjugates (Scheme 1). In this approach, photoredox-mediated single electron oxidation selectively activates the C-terminal carboxylate under visible-light irradiation with a lumiflavin photocatalyst. The short-lived carboxyl radical releases CO₂ to generate an α-amino alkyl radical, which then reacts to the Michael acceptor of the NB-alkyne linker, thereby installing a bioorthogonal alkyne handle at the peptide terminus^35,36^. Omniligase ligation, as previously described, can be directly implemented after photoredox modification of the peptide resulting in a DNA-linker-peptide-linker intermediate. This construct can be directly used for a CuAAC reaction to couple azide-dT30-desthiobiotin construct as a threading tail using CuSO_4_/THPTA complex with sodium ascorbate under mild reaction conditions^37^. The final DLPLD conjugate can be effectively purified and enriched using streptavidin-coated magnetic beads, which selectively capture the desthiobiotin-labeled products while removing unreacted DNA strands that could hinder nanopore sequencing. This workflow proceeds in aqueous conditions and is broadly compatible across very different peptide sequences.

**Scheme 1.**
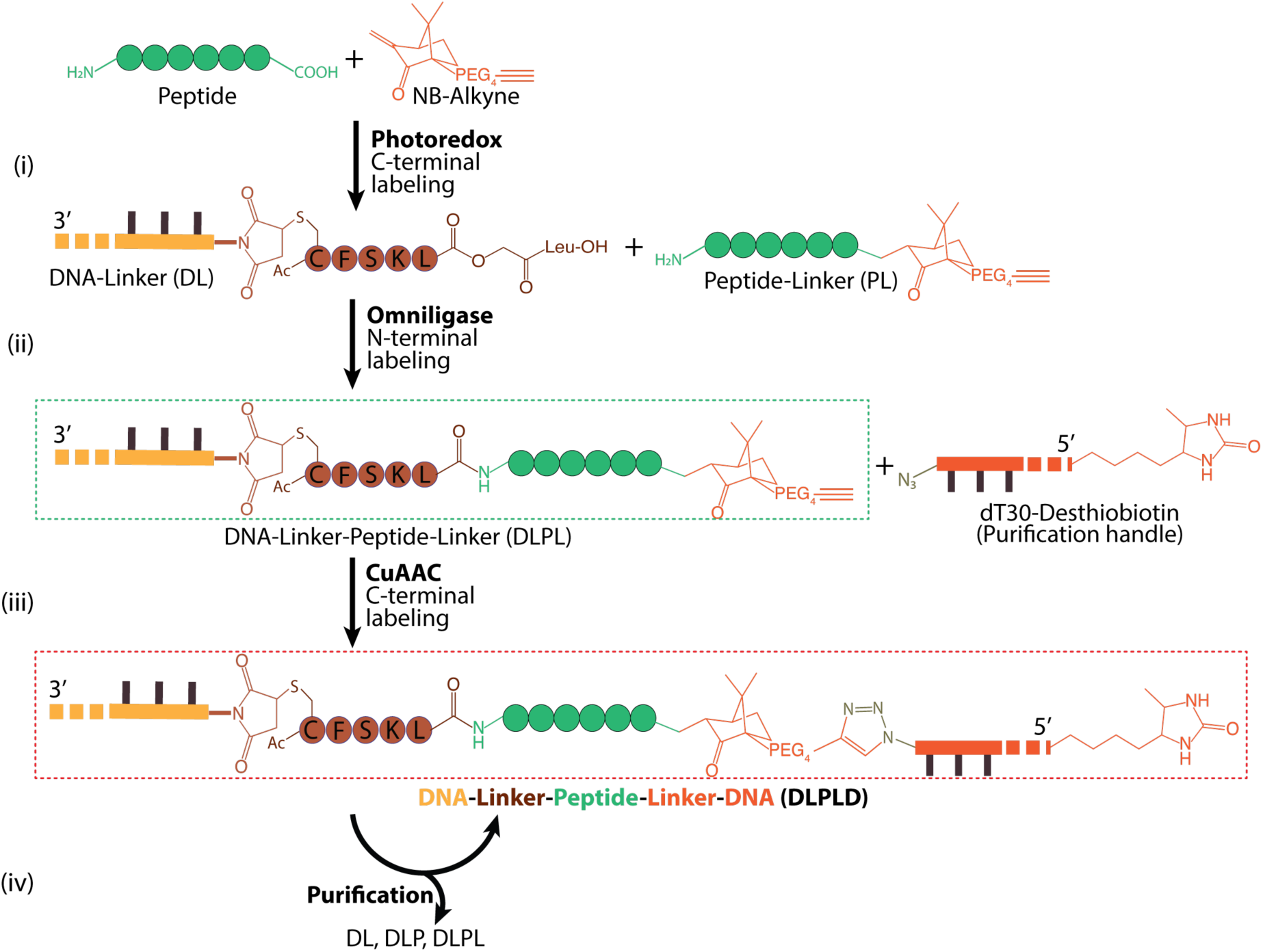
Combined N- and C-terminal conjugation strategy for constructing DNA–peptide–DNA conjugates. (i) Photoredox-mediated decarboxylative C-terminal modification installs an NB–alkyne linker on the peptide of interest to yield a peptide–linker (PL) intermediate. (ii) N-terminal attachment of a DNA–linker (DL) via omniligase ligation affords a DNA–linker–peptide–linker (DLPL) construct. (iii) In the last step, the C-terminal alkyne is coupled to an azide-functionalized dT₃₀ threading tail by CuAAC, producing the final DNA–linker–peptide–linker–DNA (DLPLD) conjugate. (iv) The dT₃₀ threading tail with a desthiobiotin purification handle enables affinity purification with streptavidin-coated magnetic beads, thereby enriching DLPLD conjugates from unreacted intermediates.

As a proof of concept, we first applied this strategy to peptides that were validated using single-terminal conjugation. Formation of DLPLD conjugates was verified by urea-PAGE, which showed distinct bands corresponding to each intermediate and the final product (Figure 3A). With each added molecular component, the conjugate molecule showed lower electro-mobility during urea-PAGE (Figure 3A). After the full conjugation workflow, peptides carried a segment of DNA on both the N-terminus and C-terminus and showed a distinct band of the full conjugate (DLPLD).

**Figure 3.**
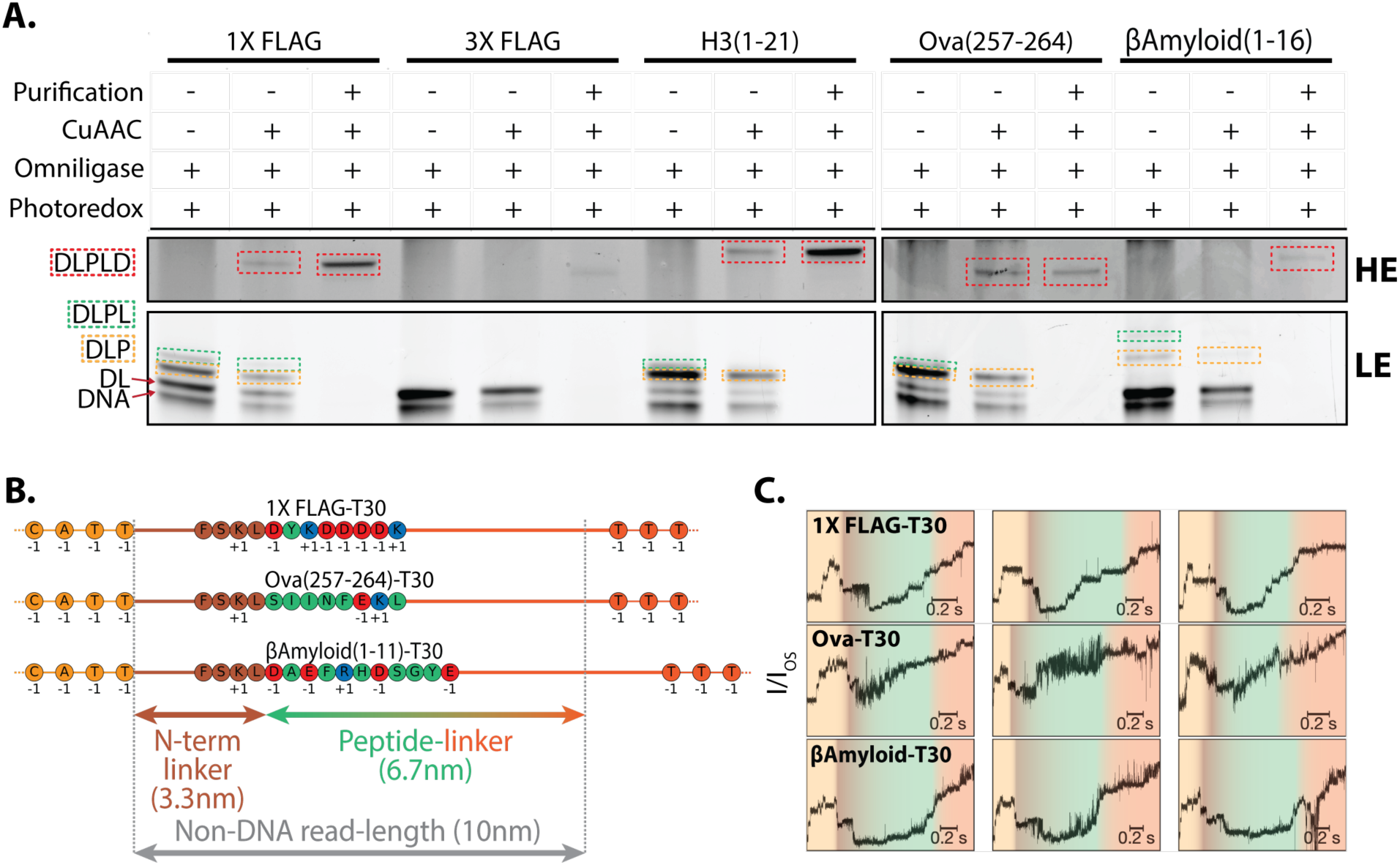
C-terminal conjugation enables threading tail attachment that is essential for sequencing short peptides. (A) Urea-PAGE (10%) analysis of DLPLD formation with various peptides. Distinct band shifts confirmed successive conjugation steps, high-exposure (HE) and low-exposure (LE) regions. Each peptide was analyzed in intermediate steps: after photoredox and omniligase ligation (Lane 1), after CuAAC reaction (Lane 2), and after affinity purification with streptavidin (Lane 3). Rectangles highlight reaction products: DLP (yellow), DLPL (green), and DLPLD (red). (B) Linear schematic of DLPLD constructs of varying length. Depending on peptide length and charge profile, a fraction of the threading tail (orange) can enter the MspA before the sequencing event ends. (C) Representative nanopore timetraces of short-peptide DLPLD constructs with a threading tail. Gradient colors indicate transitions from DNA (light yellow) to linker (brown), peptide (green), and threading tail (orange) regions. Ion currents (I) are normalized to the local unblocked open-state pore current (I_OS_). All plots are shown in 0.3 – 0.65 I/I_OS_ range.

We then proceeded to sequence the DLPLD constructs of short peptides with nanopores (Figure 3B). With a negatively charged DNA segment as the threading tail, the short peptides – which earlier failed in the DLP sequencing approach – acquired better stretching during translocation, allowing prolonged measurements (Figure 3C). Representative nanopore traces of short-peptide DLPLD constructs revealed distinct signals of the peptides that varied for different sequences. Both 1X FLAG and βAmyloid(1–11) constructs generated reproducible signatures under these conditions. These findings indicate that C-terminal threading tails enables capture and stable readout of short peptides.

However, Ova(257-264), with a net neutral charge profile, generated more variable signals. This behavior is consistent with a weaker electrophoretic stretching inside the pore, which likely permits greater conformational freedom of the peptide backbone, yielding broader current distributions, a phenomenon also observed in a recent study^19^. While weaker stretching can thus introduce variability, a previous study reported sequence-specific differences that remained discernible with a very high precision^17^, indicating that sequencing can likely still resolve peptide variants even under reduced electrophoretic tension when a threading tail is installed in the DLPLD.

We observed that most peptides tolerated photoredox modification without any loss of ligation efficiency in the subsequent omniligase reaction (Supplementary Figure 9). Thus, omniligase remained active in the presence of photoredox reaction components, allowing the two steps to be performed sequentially without intermediate purification. While this strategy was effective for most peptides (8/11, see Supplementary Table 2), some peptides showed reduced yields after photoredox-mediated ligation. Control experiments indicated that the decrease arose from peptide-specific degradation in the presence of lumiflavin and 445 nm irradiation, rather than from enzyme inhibition (Supplementary Figure 10). A subset of peptides showed greater sensitivity to the degradation. In particular, the 3X FLAG peptide produced no detectable ligation product (Figure 3A), while C(EGSG)_5_EF and C(QGSG)_5_QF exhibited markedly reduced yields compared to reactions performed without photoredox treatment (Supplementary Figure 4, Supplementary Figure 9).

### Charge neutralization enables nanopore sequencing of positively charged peptides

Positively charged peptides such as C(KGSG)_5_KF and H3(1-21) histone peptide conjugates remained among the most challenging substrates for nanopore sequencing. This is not surprising since they are subjected to an upward electrophoretic force on the cationic residues, which oppose the downward forces on DNA and negatively charged peptides. Even when equipped with threading tails, we found that these peptides failed to translocate. To overcome this limitation, we used NHS ester to acetylate the lysine side chains of C(KGSG)_5_KF construct, thereby removing the net positive charge (Figure 4A). Formation of such chemically altered DLPLD conjugates was verified by urea-PAGE, which showed distinct gel bands corresponding to each reaction step. The clean band shift after NHS-mediated acetylation, with no detectable initial product, indicated highly efficient labeling of lysine residues (Figure 4B).

**Figure 4.**
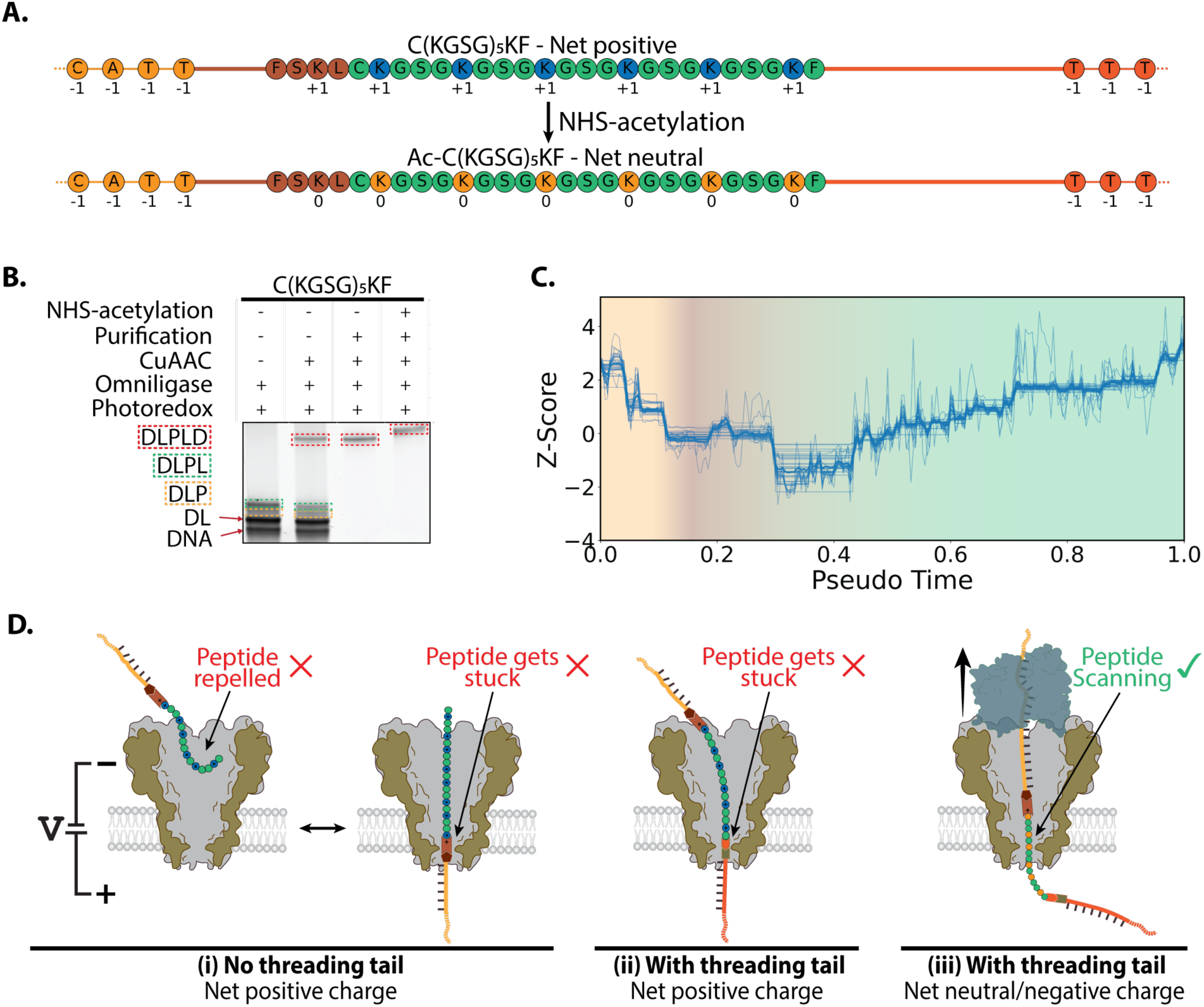
Chemical charge neutralization enables nanopore sequencing of positively charged peptides. (A) Linear schematic of C(KGSG)₅KF peptide before and after neutralization by NHS ester acetylation of lysine side chains. (B) Urea-PAGE (10%) analysis of DLPLD formation of C(KGSG)_5_KF peptide. Peptide was analyzed in intermediate steps: after photoredox and omniligase ligation (Lane 1), after CuAAC reaction (Lane 2), after affinity purification with streptavidin (Lane 3), and after NHS-acetylation (Lane 4). Rectangles indicate reaction products: DLP (yellow), DLPL (green), and DLPLD (red). (C) Dynamic time warping (DTW) alignment of peptide-region timetraces for Ac-C(KGSG)_5_KF DLPLD construct (N = 38). Gradient colors indicate transition from DNA (light yellow) to linker (brown) and peptide (green) regions. (D) Schematic models of charge-dependent translocation scenarios. (i) A peptide lacking a threading tail and carrying net positive charge is unlikely to enter the pore peptide-first and may instead engage in bidirectional “tug-of-war” when entering DNA-first. (ii) A positively charged peptide with a threading tail could enter the pore but positively charged region may fail translocating to the bottom side due to bidirectional electrophoretic forces affecting the molecule. (iii) Neutral or negatively charged peptide with threading tail can translocate through the nanopore more efficiently, allowing Hel308-mediated nanopore sequencing.

Attempts of nanopore sequencing of acetylated and non-acetylated constructs demonstrated the critical role of charge neutralization. After treating C(KGSG)_5_KF peptide with sulfo-NHS-acetate, the DLPLD of acetylated peptide (Ac-C(KGSG)_5_KF) was successfully identified in nanopore sequencing. Dynamic time warping (DTW) signal alignment^43^ revealed reproducible peptide-region signals for Ac-C(KGSG)₅KF (Figure 4C), in contrast to absent signals observed for the non-acetylated variant. Notably, the acetylated construct displayed higher signal amplitude and increased noise within the peptide region, as similarly observed for the weakly charged Ova(257–264) DLPLD analyte, suggesting weaker electrophoretic stretching of the neutralized peptide during translocation. The findings confirm that reducing cationic charge restored the sequencing ability and provided distinct signals for previously positively charged peptide. These findings align with prior observations that opposing electrophoretic force can trap or repel opposing charge peptides^44–46^.

Figure 4D sketches the variety of charge-dependent threading of peptides through a nanopore. Positively charged peptides lacking threading tails are unable to enter peptide-first (Figure 4D(i), left). Instead, upon entering DNA-first, they typically engage in a bidirectional “tug-of-war” where the negatively charged DNA handle are subjected to a downward electrophoretic force while the positively charged peptide experiences an opposing upward force (Figure 4D(i), right)^46^. When equipped with a negatively charged threading tail, such peptides can enter the pore but subsequently experience similar opposing electrophoretic forces (Figure 4D(ii)). The downward force on the DNA and threading tail competes with the upward force on the positively charged peptide region, preventing complete translocation and causing the molecule to become trapped within the nanopore. By contrast, tailed peptides with neutral or negative charge are expected to pass sensing region, which subsequently enables Hel308 to ratchet the analyte through the sensing region for sequencing (Figure 4D(iii)). Together, these results demonstrate that chemical charge neutralization provides a practical solution for expanding nanopore sequencing to highly basic peptides. More broadly, they establish charge modulation as a generalizable design principle for adapting diverse peptide substrates to nanopore analysis.

## Discussion and conclusions

This study establishes a set of design rules for adapting natural peptides to nanopore sequencing by developing terminal conjugation strategies and evaluating the effects of peptide length and charge on nanopore sequencing. Attachment of a DNA handle at one terminus provides a general entry point for DNA motor-driven peptide sequencing. Longer and negatively charged peptides yield reproducible and discriminative ionic current patterns, whereas short peptides require additional conjugation. DNA-peptide-DNA constructs enable sequencing of short and neutral peptides by enforcing capture through a negatively charged threading tail and promoting sufficient peptide stretching. Finally, chemical neutralization of positively charged peptides that would otherwise resist nanopore translocation enables sequencing of such analytes. Together, these principles define a versatile chemical toolset for adapting diverse peptides to nanopore sequencing.

The terminal conjugation chemistries introduced in this study combine enzymatic and chemical approaches to access both peptide termini under mild and orthogonal conditions. At the N terminus, omniligase provides rapid, sequence-tolerant peptide bond formation between a DNA-linked acyl donor and the target peptide, yielding DNA–peptide conjugates. At the C terminus, photoredox decarboxylation enables site-specific activation of the terminal carboxylate, installing an alkyne linker that can be coupled to azide-modified DNA through CuAAC.

While our work demonstrates feasibility across representative peptides, it offers several opportunities for further optimization. At the N terminus, the DNA-linker intermediate is based on a pentapeptide acyl donor (Ac-CFSKL-OCamLeu) containing a positively charged residue. Introducing a negatively charged amino acids here in the linker may improve electrophoretic capture and yield more uniform translocation behavior of the analyte. However, because the last four C-terminal residues of the acyl donor critically influence ligation efficiency, such modifications must be carefully designed to avoid interfering with omniligase activity^34,40^. Alternatively, chemistries such as 2-pyridinecarboxaldehyde (2-PCA) labeling could supplement omniligase in cases of low ligation efficiency^25,27^. At the C terminus, the current linker connecting DNA with a peptide is relatively long (4.9 nm) and electrically neutral; shortening it and/or introducing negative charges may enhance electrophoretic pulling forces, resulting in more consistent signal profiles, particularly for short or weakly charged sequences. Finally, integration of controlled electro-osmotic flow from the nanopore surface may provide more uniform stretching forces inside the nanopore, minimizing sequence-dependent fluctuations and yielding more reproducible readouts across various peptide substrates^47–49^.

Compared to prior nanopore peptide sequencing approaches, which relied on synthetic peptides with pre-engineered handles, our strategy broadens the accessible sample space to unmodified peptides. The sequence tolerance of our conjugation chemistries – which do not require pre-installed reactive groups or specific recognition motifs – enables analysis of native peptides or samples after protease digestion. This represents a critical step toward applying nanopore sequencing to real proteomic samples, where peptide sequences cannot be preset. This work provides a chemical foundation for single-molecule proteomic analysis beyond synthetic model systems. Future integration with protease digestion workflows will further advance nanopore sequencing toward comprehensive, single-molecule proteome characterization.

## Materials

Materials purchased for this study were Omniligase-1 enzyme (EnzyTag), Tricine (Merck, Cat # T9784), TCEP (Merck, Cat # 646547), 10X PBS (Thermo Fisher, Cat # 70011044), Lumiflavin (Cayman Chemical, Cat #20645), Cesium formate (Thermo fisher, Cat #B21327.14), Pierce Anti-FLAG Magnetic Agarose (Thermo Fisher, Cat #A36797), DynaBeads-Streptavidin (Thermo Fisher, Cat #65001), Sulfo-NHS-Acetate (TargetMol, Cat #T16959), Mini-PROTEAN 10% TBE-Urea Gel 10 well (Bio-Rad, Cat #4566033), Novex TBE-Urea Sample Buffer (2X) (Thermo Fisher, Cat #LC6876), Oligo Clean & Concentrator Kit (Zymo Research, Cat #D4061), ZR small-RNA™ PAGE Recovery Kit (Zymo Research, Cat #R1070), SYBR-Gold (Thermo Fisher, Cat#S11494). NB-PEG_4_-Alkyne linker was custom synthesized by X-Chem. Fmoc-Phe-Wang resin was purchased from Rapp Polymere (0.77 mmol/g; Cat #H0370751321), amino acids were purchased from Merck Life Science.

## Methods

### DNA-Linker preparation

Lyophilized single-stranded DNA with a 5′-maleimide group (Template DNA) was dissolved in PBS buffer (1X, pH 7.4) and immediately mixed with Ac-CFSKL-OCamLeu linker peptide to final concentrations of 25 µM DNA and 500 µM peptide. The reaction mixture was incubated for 1 h at 25 °C. Unreacted maleimide groups were quenched with 10 mM dithiothreitol (DTT) for 20 min at 25 °C. Excess unreacted linker peptide was removed using Oligo Clean & Concentrator kit columns (Zymo Research), following the manufacturer’s protocol. Columns were washed twice with 600 µL wash buffer, and DNA–linker conjugates were eluted in Milli-Q water. Product concentrations were determined by UV absorbance (NanoDrop), and aliquots were adjusted to 50 µM and stored at −20 °C.

### Omniligase ligation

Ligation reactions were carried out with either purified peptides (resuspended in Milli-Q water) or crude peptides obtained directly after photoredox modification without further purification. Reaction mixtures contained 10 µM DNA–linker, 500 µM peptide, 200 mM tricine buffer (pH 8.2), and 1 mM tris(2-carboxyethyl)phosphine (TCEP). Components were premixed on ice, and Omniligase-1 was added last to a final concentration of 10 µg/mL. Reactions were incubated for 30–60 min at 25 °C. Excess peptide and other small-molecule components were removed using Oligo Clean & Concentrator kit columns (Zymo Research) according to the manufacturer’s protocol. Columns were washed twice with 600 µL wash buffer, and DNA–linker–peptide conjugates were eluted in Milli-Q water at the desired concentration.

### PAGE purification

DNA–linker–peptide (DLP) constructs were purified using the ZR Small-RNA™ PAGE Recovery Kit (Zymo Research) following the manufacturer’s instructions. Purified samples were eluted in Milli-Q and used for nanopore sequencing experiments directly or stored at −20 °C until further use.

### Photoredox reaction

Photoredox reactions were carried out under oxygen-free conditions. DMSO, Milli-Q water, and cesium formate buffer (10 mM, pH 3.5) were degassed by sparging with nitrogen for 20 min, and all further steps were performed in a nitrogen glove box. Lyophilized peptides were dissolved in degassed Milli-Q water to a concentration of 2 mM. Reaction mixtures contained 25 µM lumiflavin, 25 mM NB–PEG₄–alkyne linker, 10 mM cesium formate, 1 mM peptide, and 5% (v/v) DMSO. Samples were transferred into tightly sealed HPLC glass vials and irradiated with 445 nm blue LEDs (Lumidox II, 115 mW, active cooling fan, 30 °C) for 6 h. Lumiflavin was replenished every 2 h with 50% of the initial concentration. After irradiation, peptide products were either used directly for Omniligase ligation or stored at −20 °C until further use.

### CuAAC reaction

Samples obtained after photoredox and Omniligase reactions were used for DNA tail attachment via CuAAC chemistry. CuSO₄ (10 mM) and THPTA ligand (20 mM) were premixed in Milli-Q water 5 min prior to the reaction. Sodium ascorbate was freshly prepared in Milli-Q water. Reaction mixtures contained ∼1 µM purified DNA-linker-peptide-linker (DLPL) conjugate, 50 µM DTB–T₃₀–azide DNA, 0.5 mM CuSO₄, 1 mM THPTA, 2 mM Sodium ascorbate, and 1X PBS buffer (pH 7.4). Reactions were incubated for 2 h at 25 °C. Non-DNA components were removed using Oligo Clean & Concentrator kit columns (Zymo Research) according to the manufacturer’s protocol. Columns were washed once with 600 µL wash buffer, and products were eluted in Milli-Q water at the desired concentration.

### Desthiobiotin-Streptavidin affinity purification

Products containing the threading tail were purified from the unreacted components: DNA, DNA-linker, DNA-linker-peptide, DNA-linker-peptide-linker. Using a desthiobiotin handle available on 5’ of the threading tail and DynaBeads-Streptavidin, final product of DNA-linker-peptide-linker-peptide was purified. Notably, a large fraction of unreacted DTB-T_30_-Azide DNA still remained in the sample, and consequently the binding capacity was mainly accounted to the DTB-T_30_-Azide molecule.

DynaBeads-Streptavidin (100 µL per 200 pmol desthiobiotin) were washed three times with buffer 2X BW buffer, then resuspended in 1X binding buffer together with the sample and incubated with beads for 30 min at room temperature with orbital rotation. After binding, beads were separated with a magnet and washed 4 times with 1X PBS (pH 7.4) + 0.05% Tween-20. Bound material was eluted by competition with 5mM biotin in 1X PBS for 10 min at 37 °C, to displace bound conjugates. Elution was repeated twice and collected material was desalted using Oligo Clean & Concentrator kit columns (Zymo Research) according to the manufacturer’s protocol. Columns were washed once with 600 µL wash buffer, and products were eluted in Milli-Q water at the desired concentration. Purified samples were analyzed by 10% denaturing Urea-PAGE.

### Urea-PAGE

Samples containing DNA were analyzed on 10% denaturing polyacrylamide gels containing 7 M urea, run in 1X TBE buffer. Gels were prerun at 200 V for 30 min prior to loading. Samples were premixed with 1X sample loading buffer and 5 µL of the sample was loaded directly. Electrophoresis was performed at 200 V for 45 min. After electrophoresis, gels were rinsed twice with 50 mL Milli-Q water and stained with 5mL 1X SYBR Gold in Milli-Q for 3 min with gentle agitation. Gels were subsequently rinsed twice with 50 mL Milli-Q water before imaging. Fluorescent images were acquired using a Typhoon scanner (Cytiva) at 50 µm resolution, with excitation at 488 nm and emission detection through a 525 nm ± 10 nm band-pass (BP20) filter.

### Nanopore sequencing

Before the sequencing experiments, DLP or DLPLD samples were annealed with Complementary DNA containing cholesterol tag on 3’ end (Supplementary Table 2). Samples were mixed to a final 500 nM concentration in sample buffer (500 mM KCl, 10mM HEPES, pH 8.0). Annealing program of 80 °C (5 min) → −0.2 °C/s → 20 °C was used for all samples for DNA hybridization.

Nanopore experiments were performed using MinION platform with pre-inserted MspA-M2(N93D) nanopore flow cells provided by Oxford Nanopore Technologies. Run conditions were set with a custom software (available from Oxford Nanopore Technologies). The experiments were performed at 36 °C and a constant voltage of −180 mV with a 5 kHz sampling frequency and periodic voltage flips every 30 s.

Before each sequencing run, flow cells were primed via the priming port with 800 µL wash buffer (500 mM KCl, 10 mM MgCl_2_, 10 mM HEPES, pH 8.0) and incubated for 5 min. Sequencing buffer (200 µL; 500 mM KCl, 10 mM MgCl_2_, 10 mM HEPES, 1 mM ATP, pH 8.0) was then added to the priming port. A 75 µL of sample was loaded onto the SpotON port containing 5 nM annealed DLP or DLPLD and 1 µM Hel308 in sequencing buffer (500 mM KCl, 10 mM MgCl_2_, 10 mM HEPES, 1 mM ATP, pH 8.0).

### Nanopore sequencing data analysis

With ONT’s fast5_research tools (https://github.com/nanoporetech/fast5_research), the sequencing data files were processed with a custom multi-threaded python pipeline. First, open-state current levels for each channel (1-512) were determined by fitting a Gaussian mixture model (n=3) to ionic current distributions. Events were then extracted by segmenting regions between open states, considering only data within the 175-185 mV sequencing voltage range. Current step transitions were detected using a sensitivity threshold of 3 with minimum step length of 10 data points with the same algorithm described in previous work^1^. Raw events were filtered by satisfying minimum duration (≥1 s), step count threshold (>45 steps). For each filtered event. Events from each channel passing all quality filters were compiled into a collection and went through DNA-peptide boundary detection. The boundary detection was achieved by adapting the alignment algorithm (explained in previous work^1^) to match moving window segments of the nanopore signals to the predicted DNA signature. With a best match, the end of the signal window is selected as the DNA-peptide boundary, producing a consistent and reproducible segmentation for all events. The events with marked boundaries then went through further filtering to remove peptide signal outliers and the collection of peptide signals were used for generating signals consensus.

The peptide signal consensus was generated by a barycenter averaging algorithm with MATLAB’s built-in dynamic time warping (DTW) function. Briefly, the template for the barycenter is chosen by finding the signal trace that has the total minimal DTW distance to all other traces within the collection. Then all other traces are DTW-aligned to the template, and the barycenter average is taken by averaging the values from all traces at each point.

### NHS-ester-mediated lysine acetylation

Lysine acetylation was carried out using sulfo-NHS-acetate. C(KGSG)_5_KF sample after streptavidin purification was incubated in 100 mM NaHCO₃ buffer (pH 8.3) with 1 mM sulfo-NHS-acetate, freshly prepared in NaHCO₃ buffer to minimize hydrolysis. Reactions proceeded for 20 min at room temperature. Excess reagent was removed by purification using Oligo Clean & Concentrator columns (Zymo Research) according to the manufacturer’s protocol, and DNA-peptide conjugates were eluted in Milli-Q water for downstream applications.

### IAA capping

Cysteine containing peptides that were used for photoredox reaction were alkylated with iodoacetamide (IAA) to prevent from competitive thiol-conjugate addition reported in previous study^2^. For this reaction freshly made 5 mM iodoacetamide (IAA) was mixed with 1 mM peptide in 100 mM HEPES pH 8.0 and incubated for 20 min at RT in the dark. Excess of IAA was quenched using 50 mM DTT for 10 min. Alkylated peptides were purified on an AdvanceBio Peptide Map C18 column (4.6 × 150 mm, 2.7 µm, Agilent) using an Agilent 1260 Infinity II HPLC system at 60 °C. Mobile phase A was water with 0.2% trifluoracetic acid and mobile phase B was acetonitrile with 0.2% trifluoracetic acid. Elution was carried out at 0.70 mL/min with a linear gradient from 5 to 40% B over 20 min, followed by a wash at 95% B and re-equilibration at 5% B. Elution was monitored at 214 nm and 260 nm, fractions corresponding to the target peak were collected and lyophilized.

### 1X FLAG and 3X FLAG affinity purification using Anti-FLAG resin

For DLPs containing FLAG peptides, magnetic agarose beads were applied directly after the Omniligase reaction. Anti-FLAG magnetic agarose beads were equilibrated with washing buffer (1X PBS, 0.2% Tween-20) by three wash cycles prior to use and then incubated with the DLP products overnight at 4 °C under gentle rotation. After incubation, beads were washed five times with washing buffer to remove unbound material.

For elution, beads were resuspended in 3X FLAG peptide (1.5 mg/mL) prepared in diluted washing buffer (0.2X PBS, 0.04% Tween-20) and incubated for 2 h at 4 °C under gentle rotation. The supernatant was collected following magnetic separation, and residual material was recovered by an additional wash with Milli-Q water. Excess FLAG peptide used for elution was removed by purification with Oligo Clean & Concentrator columns (Zymo Research) according to the manufacturer’s protocol, and purified DLPs were eluted in Milli-Q water for downstream applications.

### Peptide synthesis

PSK Peptides were synthesized following Fmoc/*t*Bu Solid-Phase Peptide Synthesis (SPPS) strategy using an automated peptide synthesizer (CS336X Peptide Synthesizer CS BIO Co.). In general, amino acids were added as follows: the resin was pre-swollen with dichloromethane (DCM). The Fmoc-protecting group was removed using 20% piperidine in *N,N*-dimethylformamide (DMF, 2 × 8 min). The resin was then washed with DMF (3 × 2 min). The next amino acid was treated with 2-(1H-benzotriazole-1-yl)-1,1,3,3-tetramethyluronium hexafluorophosphate (HBTU), 1-hydroxybenzotriazole (HOBt), and *N,N*-diisopropylethylamine (DIPEA) in DMF for 2 min before being added to the resin. The reaction mixture was allowed to couple for 2 h, and reaction completion was monitored by resin staining using ninhydrin (15 g/L, supplemented with 30 mL/L acetic acid in *n*-butanol). Upon reaction completion, the resin was washed with DMF (3 × 2 min), deprotected by 20% piperidine in DMF (2 × 8 min), and washed again with DMF (3 × 2 min). The same steps were repeated until the desired sequence was obtained.

Chain elongation was initiated from Fmoc-Phe-Wang resin 100-200 mesh Novabiochem®, a *p*-alkoxybenzyl alcohol polymer-bound (copolystyrene-1% DVB) amino acid (loading capacity 0.77 mmol/g). chain elongation was performed with Fmoc-Gly-OH, Fmoc-Ser(*t*Bu)-OH, Fmoc-Glu(O*t*Bu)-OH, Fmoc-Gln(Trt)-OH, or Fmoc-Lys(Boc)-OH, depending on the peptide and position in the sequence. The standard protocol relied on a coupling time for each amino acid of 3h at room temperature, under continuous shaking.

After completion of the sequence by SPPS, the peptides were cleaved from the resin by treatment with a cocktail of 90% trifluoroacetic acid (TFA), 2.5% triisopropylsilane (TIS), and 7.5% Milli-Q for 2 h (10 mL TFA cocktail/1 g initial resin). The cleaved peptides were precipitated by dropwise addition of the TFA cocktail-peptide mixture to ice-cold diethyl ether (1:1 ether:hexane, 10× initial cocktail volume) and the cleaved peptide resin was washed once with a small amount of fresh cleavage cocktail. The precipitate was centrifuged for 10 min at 6000 rpm. The supernatant was discarded and the precipitate was washed with ice-cold diethyl ether and again centrifuged for 10 min at 6000 rpm. This washing step was repeated once more. The resulting precipitate was then dried in a light stream of N_2_, redissolved in MeCN:Milli-Q (1:1), snap freezed with liquid N_2_ and then lyophilized (Labconco FreeZone lyophilizer, 2.5 L, –84 °C, connected to a 35_i_ xDS Edwards Oil-Free Dry Scroll Pump).The obtained peptides were purified by semi-preparative reverse phase HPLC (Agilent 1260 Preparative HPLC with a DAD G7115A and MSD) using a preparative semi-preparative Zorbax Eclipse column (XDB-C18, 9.4 x 250 mm, 5-Micron). The eluent for purification contained for buffer A: MilliQ:MeCN – 95:5 (v/v, 0.1% TFA), and for buffer B: MilliQ:MeCN – 5:95 (v/v, 0.1% TFA), which were applied with the following gradient: 100% A for 5 min → 0%A in 20 min → 0% A for 5 min → 100% A in 5 min → 100% A for 5 min (total runtime: 40 min). The purified peptides were then lyophilized for further analysis and use.

### HPLC purification

Obtained deprotected peptides were purified by semi-preparative reverse phase HPLC (Agilent 1260 Preparative HPLC with a DAD G7115A and MSD) using a preparative semi-preparative Zorbax Eclipse column (XDB-C18, 9.4 x 250 mm, 5-Micron). The eluent for purification contained for buffer A: MilliQ:MeCN – 95:5 (v/v, 0.1% TFA), and for buffer B: MilliQ:MeCN – 5:95 (v/v, 0.1% TFA), which were applied with the following gradient: 100% A for 5 min → 0%A in 20 min → 0% A for 5 min → 100% A in 5 min → 100% A for 5 min (total runtime: 40 min). The obtained purified peptides were then lyophilized for further analysis and use.

## Data and code availability

The raw data and analysis code will be deposited in Zenodo.

## Supporting Information

Experimental procedures, additional supporting information is available from the author.

## Author Information

Corresponding Author:

Cees Dekker − Department of Bionanoscience, Kavli Institute of Nanoscience Delft, Delft University of Technology, Delft, 2629 HZ, The Netherlands;

Authors:

Justas Ritmejeris − Department of Bionanoscience, Kavli Institute of Nanoscience Delft, Delft University of Technology, Delft 2629 HZ, The Netherlands

Xiuqi Chen − Department of Bionanoscience, Kavli Institute of Nanoscience Delft, Delft University of Technology, Delft 2629 HZ, The Netherlands

Bauke Albada − Laboratory of Organic Chemistry, Wageningen University & Research, Wageningen 6807 WE, The Netherlands

## Author contributions

J.R. performed peptide conjugation and nanopore sequencing experiments, analyzed nanopore data, and drafted the manuscript. X.C. developed nanopore data analysis tools, performed nanopore data analysis, generated experimental materials, and contributed to project ideation. B.A. provided project supervision and guidance on the chemistry, including peptide synthesis. C.D. conceived the study and provided project supervision and guidance. All authors contributed to manuscript revision.

## Acknowledgments

The authors thank R. Eelkema, P. Bohlander, and J. Swaminathan for helpful discussions and technical assistance with the photoredox reaction; A. Koijen, S. van den Beelen, and L. van den Bos for valuable input on implementing the Omniligase reaction; J. Smith, M. Jordan, K. Sabharwal, and J. Wallace for discussions and for providing MinION flowcells; J. Firet for peptide synthesis; and I. Nova, M. Filius, J. van der Sande, E. van der Sluis, J. van der Torre, L. Yu, T. Eriksson, T. Hoekstra, and B. Analikwu for helpful discussions.

## Supplementary tables

**Supplementary Table 1.** Peptides used in the study for terminal conjugation and nanopore sequencing experiments.

| Name | Sequence (N→C) | Purity | Source | Modifications |
| --- | --- | --- | --- | --- |
| Linker-peptide | CFSKL | >95% | Biosynth | N term:<br>acetylation;<br>C term:<br>O-CamLeuOH |
| 1X FLAG | DYKDDDDK | >95% | GenScript | - |
| 3X FLAG | MDYKDHDGDYKDHDIDYKDDDDK | >90% | Merck | - |
| H3(1-21) | ARTKQTARKSTGGKAPRKQLA | >95% | Eurogentec | - |
| Ova257-264 | SIINFEKL | >97% | Merck | - |
| βAmyloid1-11 | DAEFRHDSGYE | >95% | Anaspec | - |
| βAmyloid1-16 | DAEFRHDSGYEVHHQK | >95% | Anaspec | - |
| βAmyloid17-28 | LVFFAEDVGSNK | >95% | Anaspec | - |
| C(EGSG)6Y | CEGSGEGSGEGSGEGSGEGSGEGS<br>GY | >95% | Custom<br>synth. | C-term:<br>amidation |
| C(EGSG)5EF | CEGSGEGSGEGSGEGSGEGSGEF | >95% | Custom<br>synth. |  |
| C(QGSG)5QF | CQGSGQGSGQGSGQGSGQGSGQF | >95% | Custom<br>synth. |  |
| C(KGSG)5KF | CKGSGKGSGKGSGKGSGKGSGKF | >95% | Custom<br>synth. |  |

**Supplementary Table 2.** Summary of peptide modifications reactions tested with three conjugation methods. Signs indicate reaction efficiency: ++ good; + low; - no reaction product; NA not applicable reaction.

| No. | Name | Peptide length | Omniligase N-terminal | Photoredox C-terminal | CuAAC C-terminal |
| --- | --- | --- | --- | --- | --- |
| 1. | 1X FLAG | 8 | ++ | + | + |
| 2. | 3X FLAG | 23 | + | - | - |
| 3. | H3(1-21) | 21 | ++ | ++ | ++ |
| 4. | Ova257-264 | 8 | ++ | + | + |
| 5. | βAmyloid1-11 | 11 | + | + | + |
| 6. | βAmyloid1-16 | 16 | + | + | + |
| 7. | βAmyloid17-28 | 12 | + | - | - |
| 8. | C(EGSG)<sub>6</sub>Y | 26 | + | NA | NA |
| 9. | C(EGSG)<sub>5</sub>EF | 23 | ++ | + | + |
| 10. | C(QGSG)<sub>5</sub>QF | 23 | ++ | + | + |
| 11. | C(KGSG)<sub>5</sub>KF | 23 | ++ | ++ | ++ |

**Supplementary Table 3.**
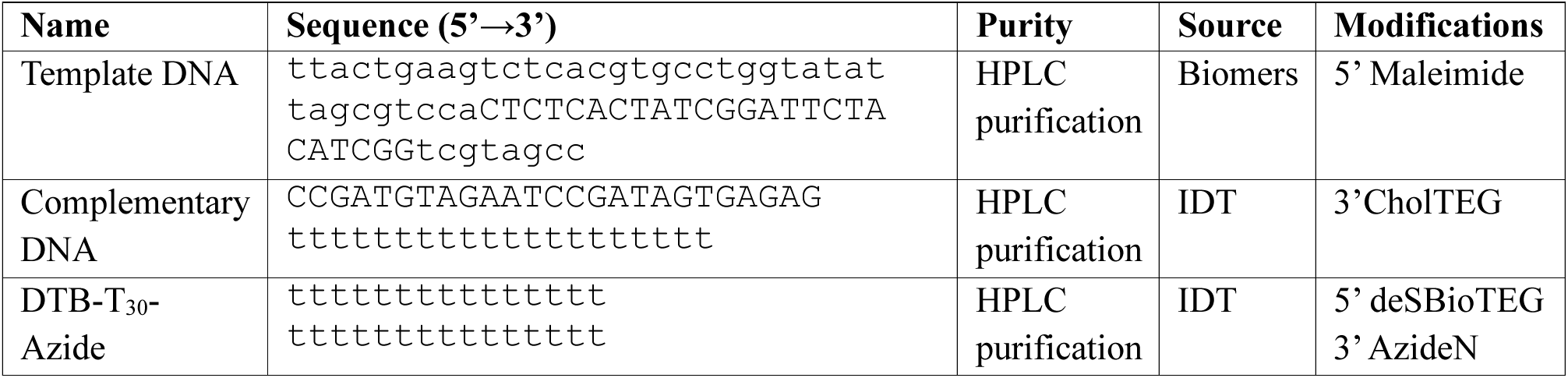
Oligonucleotides used in the study. Capital letters indicate the annealing region between the Template DNA and the Complementary DNA strand.

## Supplementary figures

**Supplementary Figure 1.**
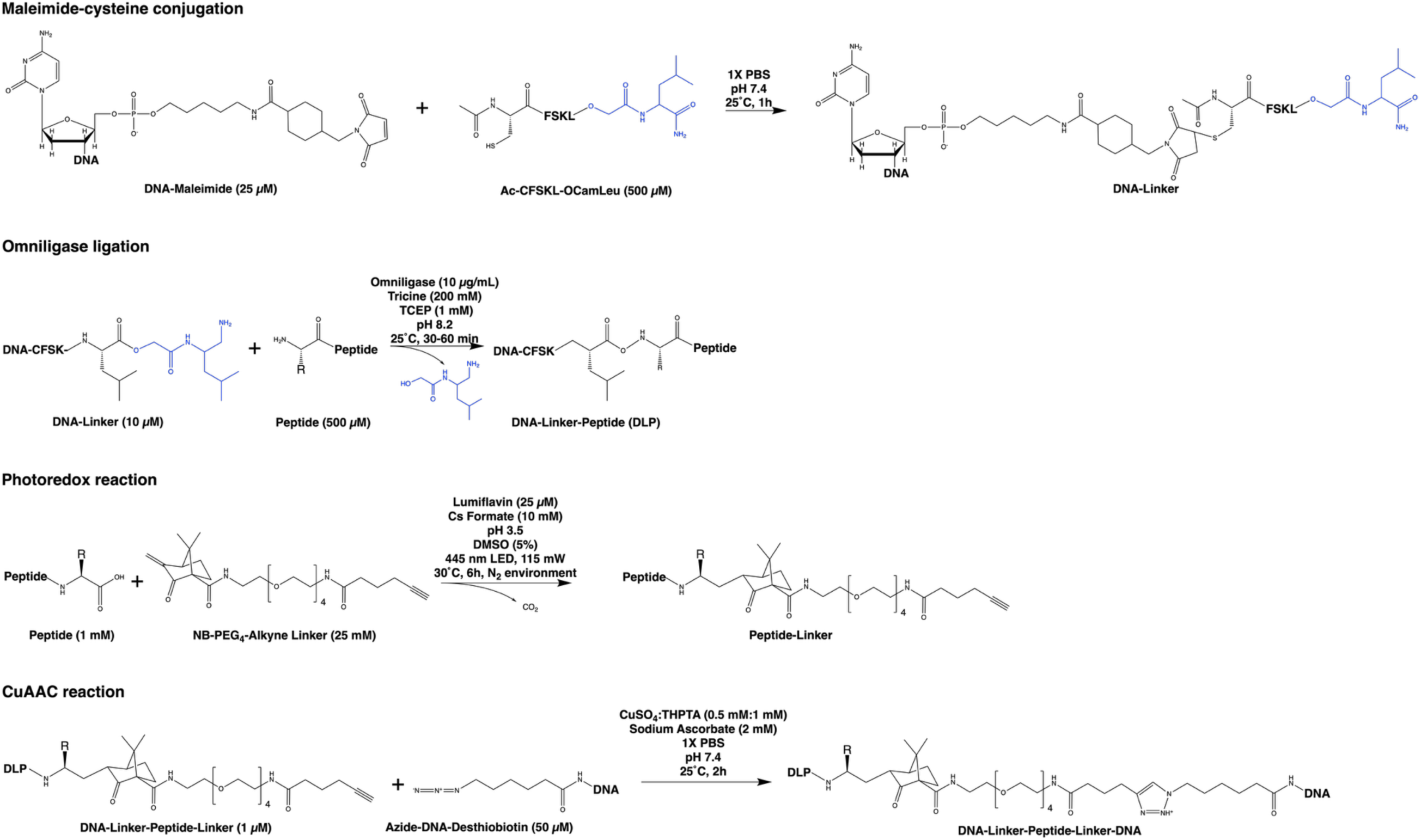
Terminal conjugation reaction schemes.

**Supplementary Figure 2.**
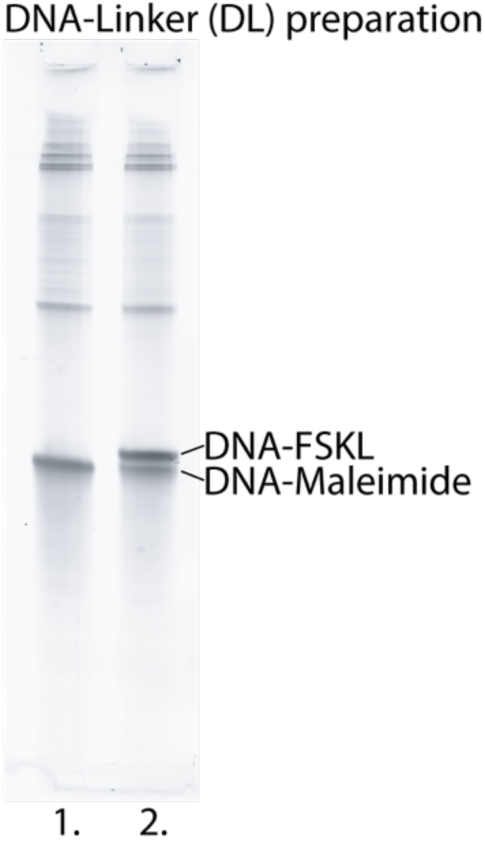
DNA–linker (DL) preparation by maleimide–cysteine coupling. Lane 1: 5′-maleimide DNA before CFSKL linker addition; lane 2: reaction after peptide addition, showing two bands: unreacted DNA–maleimide (quenched with DTT) and the DNA–FSKL product (upper band). Gel: 10% Urea-PAGE, stained with SYBR Gold.

**Supplementary Figure 3.**
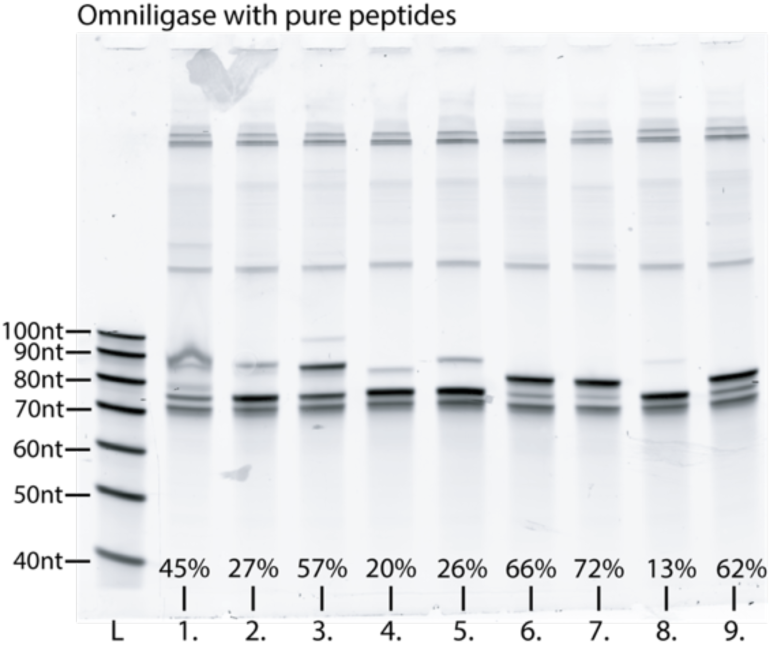
Comparison of Omniligase ligation products obtained with pure peptides. Lanes: L. 20/100 ssDNA ladder; 1. C(KGSG)₅KF; 2. C(QGSG)₅QF; 3. C(EGSG)₅EF; 4. β-Amyloid (17–28); 5. β-Amyloid (1–16); 6. Ova(257–264); 7. H3(1–21); 8. 3X FLAG; 9. 1X FLAG. Conversion yields were estimated from band intensity ratios of DLP / (DLP + DNA–FSKL) using GelGenie^3^. Gel: 10% Urea-PAGE, stained with SYBR Gold.

**Supplementary Figure 4.**
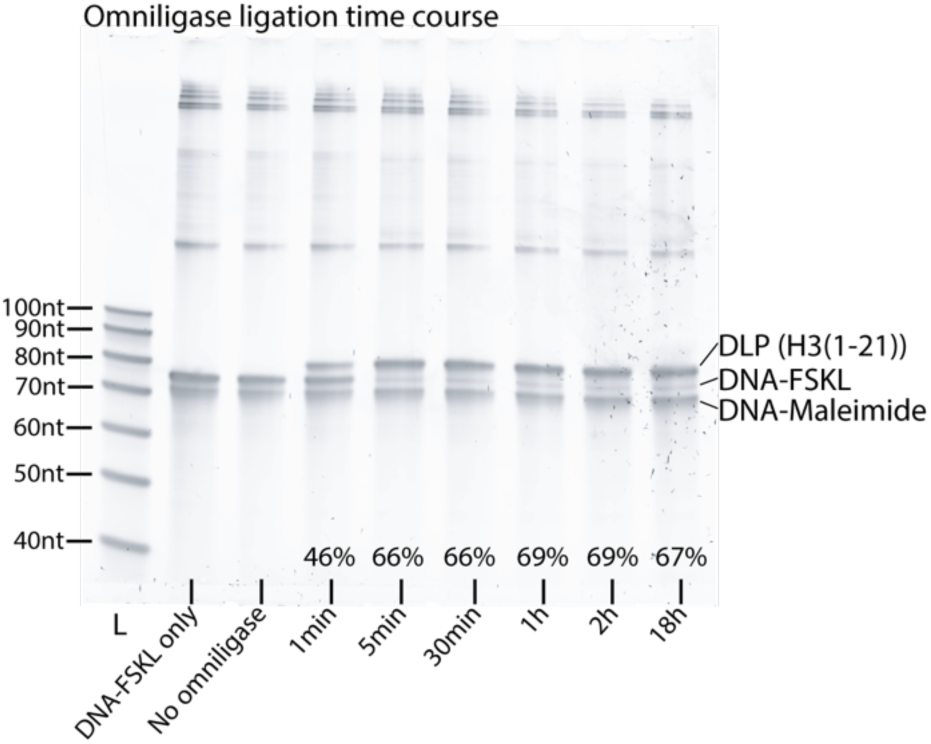
Comparison of Omniligase ligation products obtained with different incubation times using H3(1-21) peptide as a substrate. Conversion yields (shown as percentages) were estimated from band intensity ratios of DLP / (DLP + DNA–FSKL) using GelGenie^3^. L: 20/100 ssDNA ladder. Gel: 10% Urea-PAGE, stained with SYBR Gold.

**Supplementary Figure 5.**
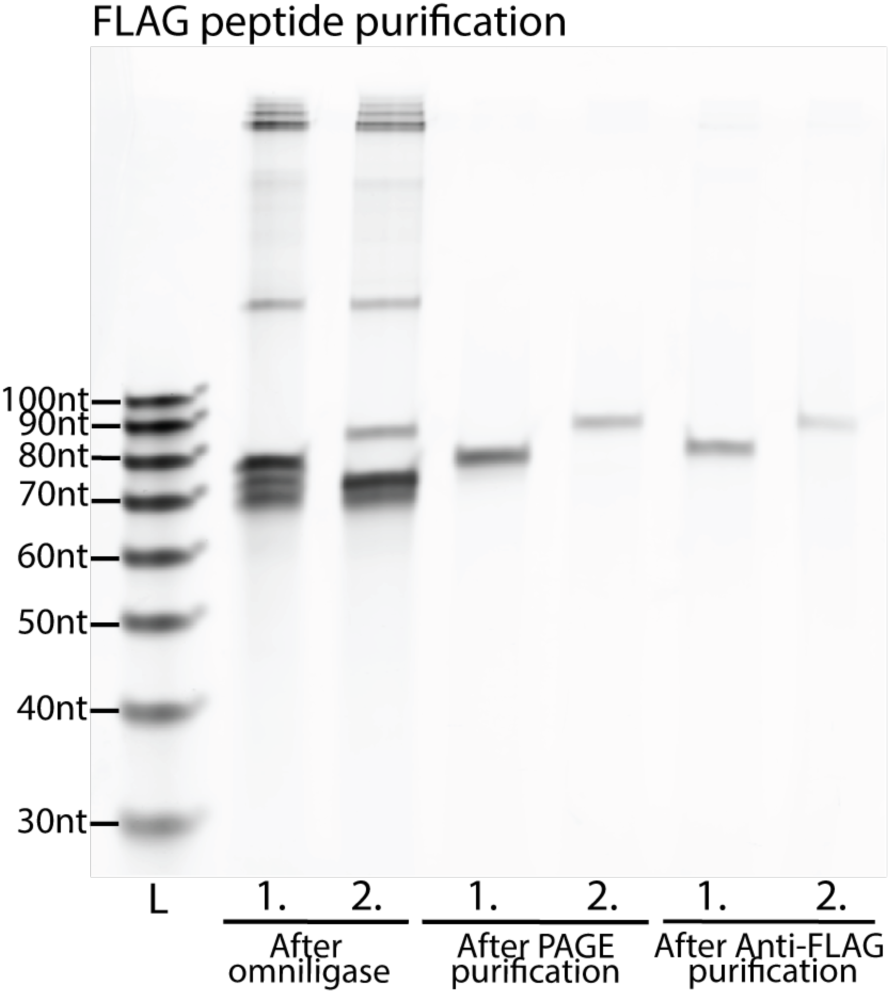
Comparison of purification methods for FLAG peptides. Lanes: 1. 1X FLAG; 2. 3X FLAG. For each peptide, samples are shown after Omniligase ligation, after PAGE purification, and after affinity purification with anti-FLAG resin. L. 20/100 ssDNA ladder. Gel: 10% Urea-PAGE, stained with SYBR Gold.

**Supplementary Figure 6.**
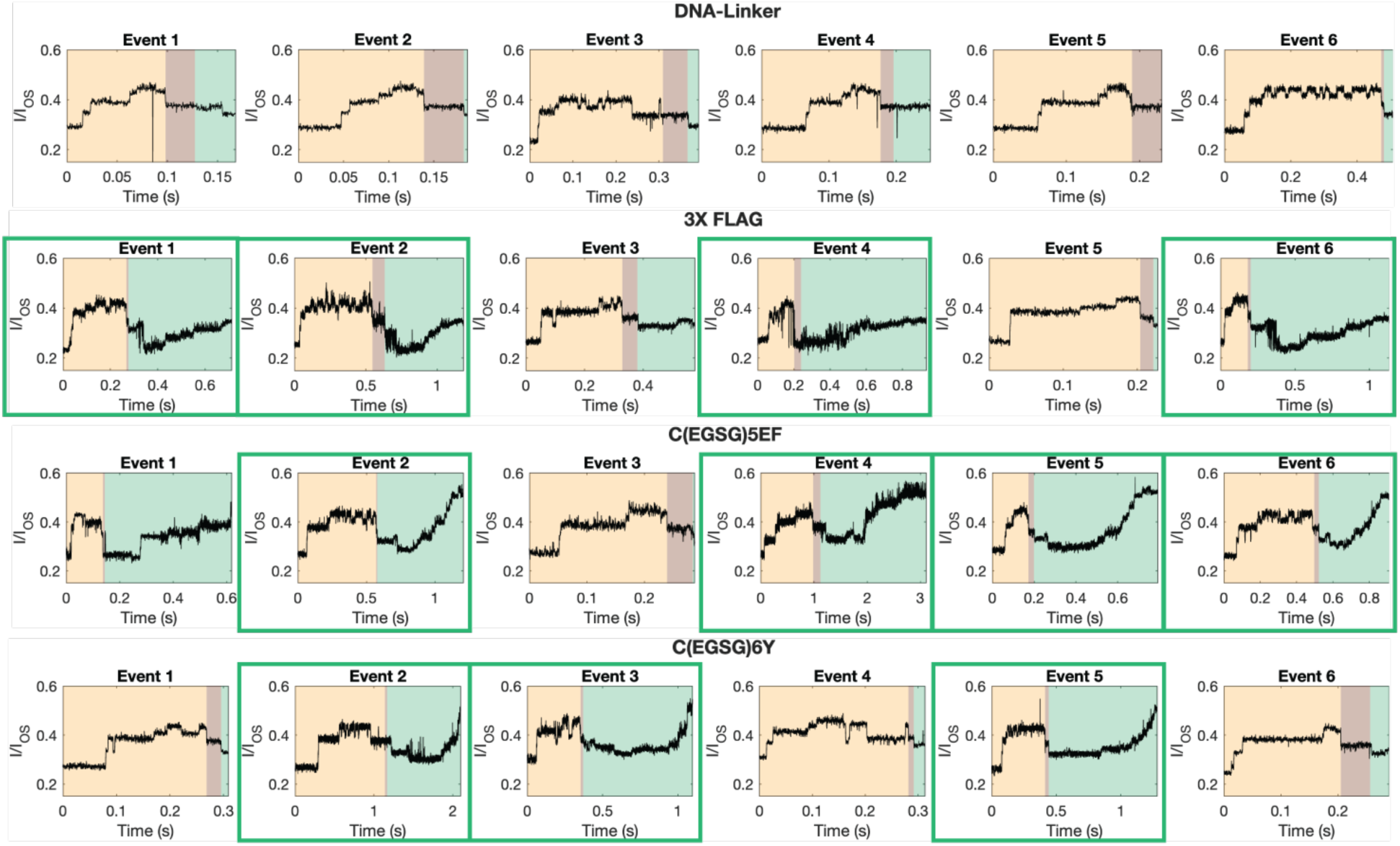
Representative sequencing time traces at the DNA–peptide junction for long-peptide constructs. Green rectangles highlight events with distinct peptide features that were used for Dynamic Time Warping alignment in Figure 2B. Events are segmented into the final DNA region (light yellow), the DNA–peptide transition (light brown), and the post-DNA peptide region (light green).

**Supplementary Figure 7.**
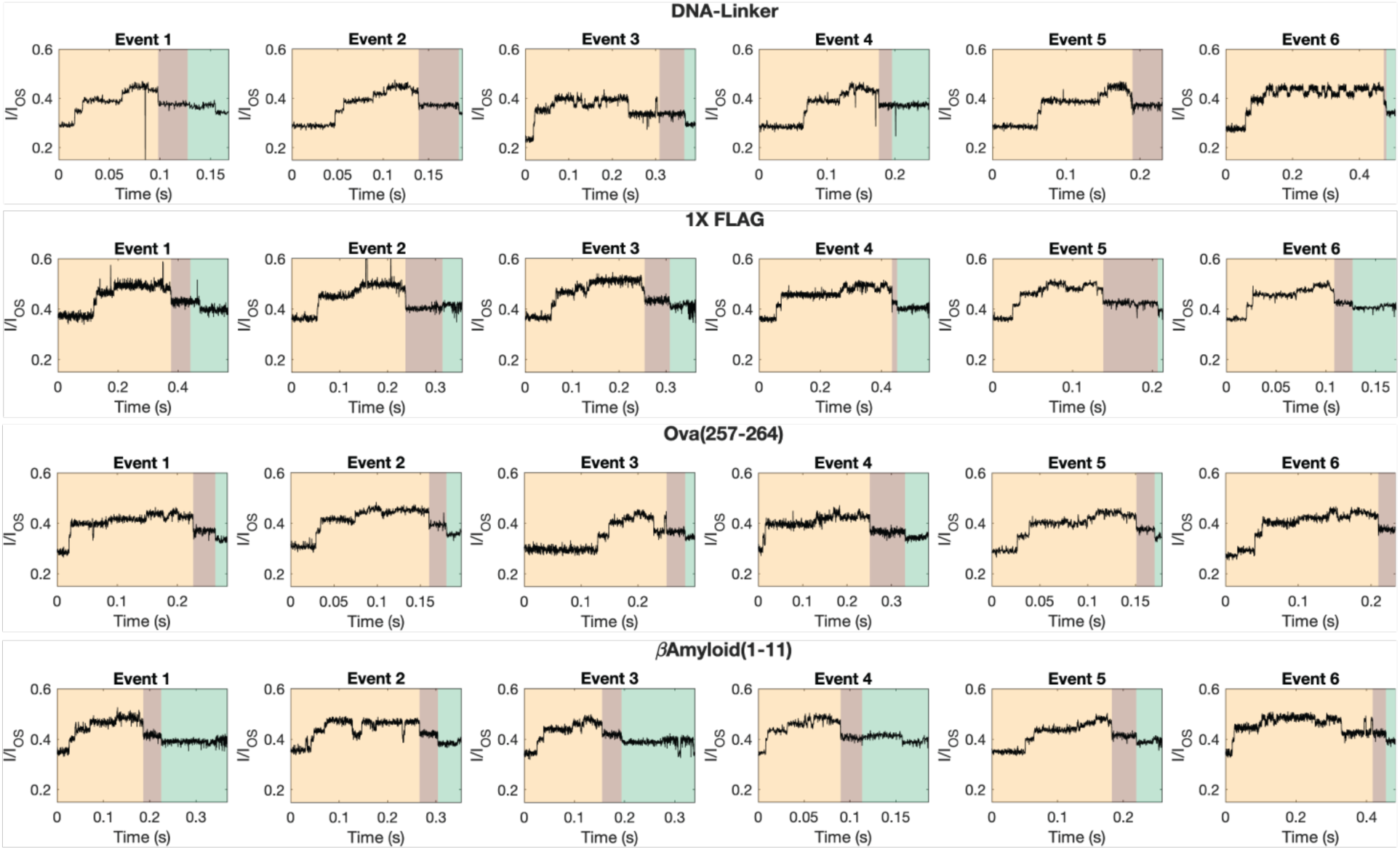
Representative sequencing timetraces at the DNA–peptide junction for short-peptide constructs. Events are segmented into the final DNA region (light yellow), the DNA–peptide transition (light brown), and the post-DNA peptide region (light green).

**Supplementary Figure 8.**
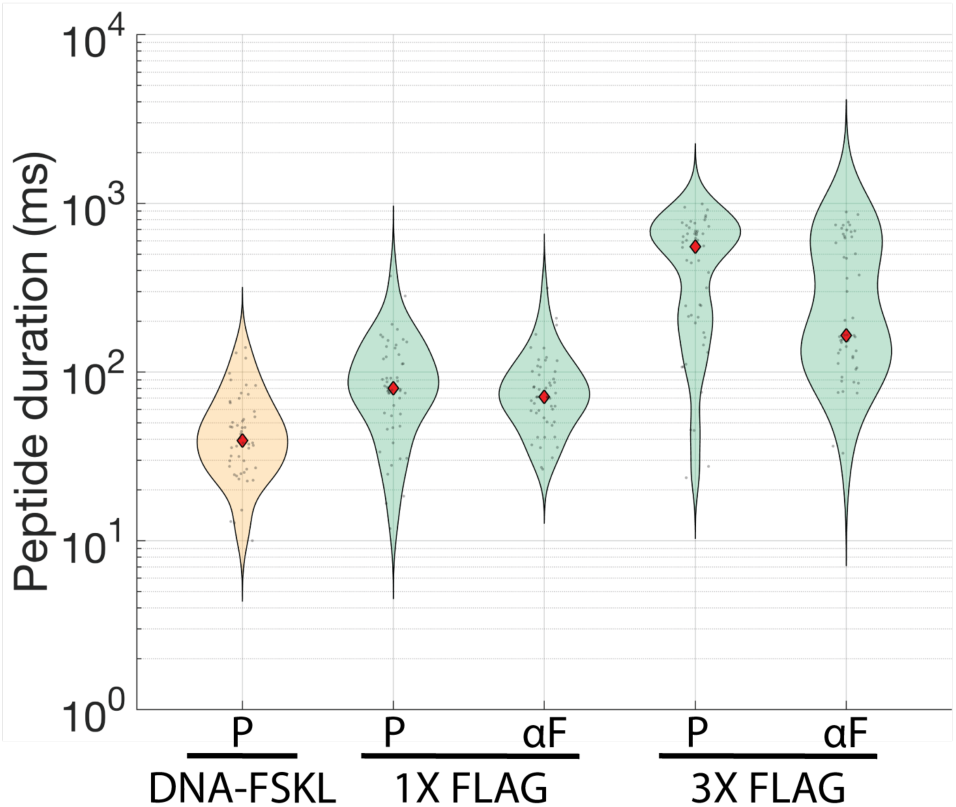
Comparison of event durations in the after-DNA region for samples purified using different methods. P indicates PAGE-purified samples; αF indicates anti-FLAG affinity–purified samples. Median event durations: DNA–FSKL (P) = 43 ms; 1X FLAG (P) = 89 ms; 1X FLAG (αF) = 73 ms; 3X FLAG (P) = 575 ms; 3X FLAG (αF) = 189 ms. For each sample, 65 randomly selected events containing a DNA signal were manually segmented as shown in Supplementary Figures 6 and 7 by combining a duration of DNA-peptide transition region and post-DNA region.

**Supplementary Figure 9.**
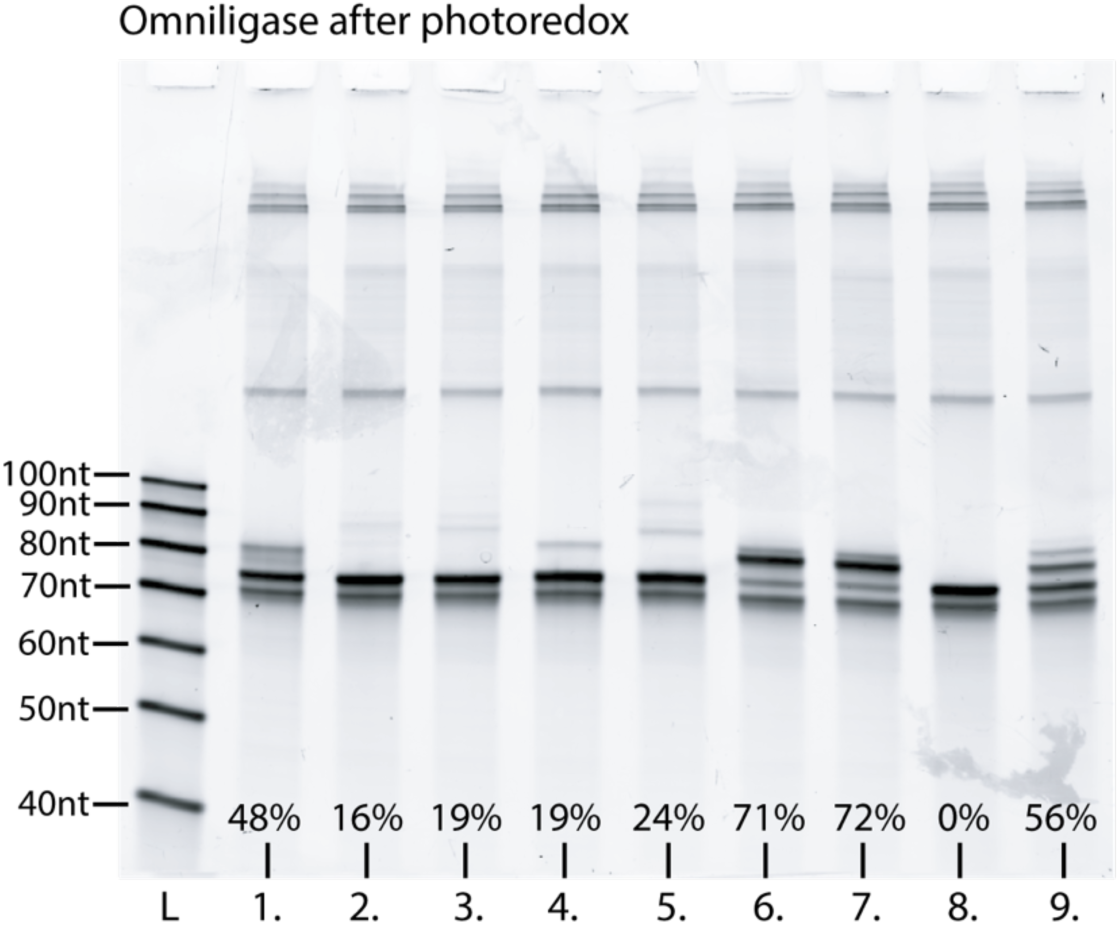
Comparison of Omniligase ligation products obtained with peptides directly after photoredox reaction. Lanes: L. 20/100 ssDNA ladder; 1. C(KGSG)₅KF; 2. C(QGSG)₅QF; 3. C(EGSG)₅EF; 4. β-Amyloid (17–28); 5. β-Amyloid (1–16); 6. Ova(257–264); 7. H3(1–21); 8. 3X FLAG; 9. 1X FLAG. Conversion yields were estimated from band intensity ratios of DLP / (DLP + DNA–FSKL) using GelGenie^3^. Gel: 10% Urea-PAGE, stained with SYBR Gold.

**Supplementary Figure 10.**
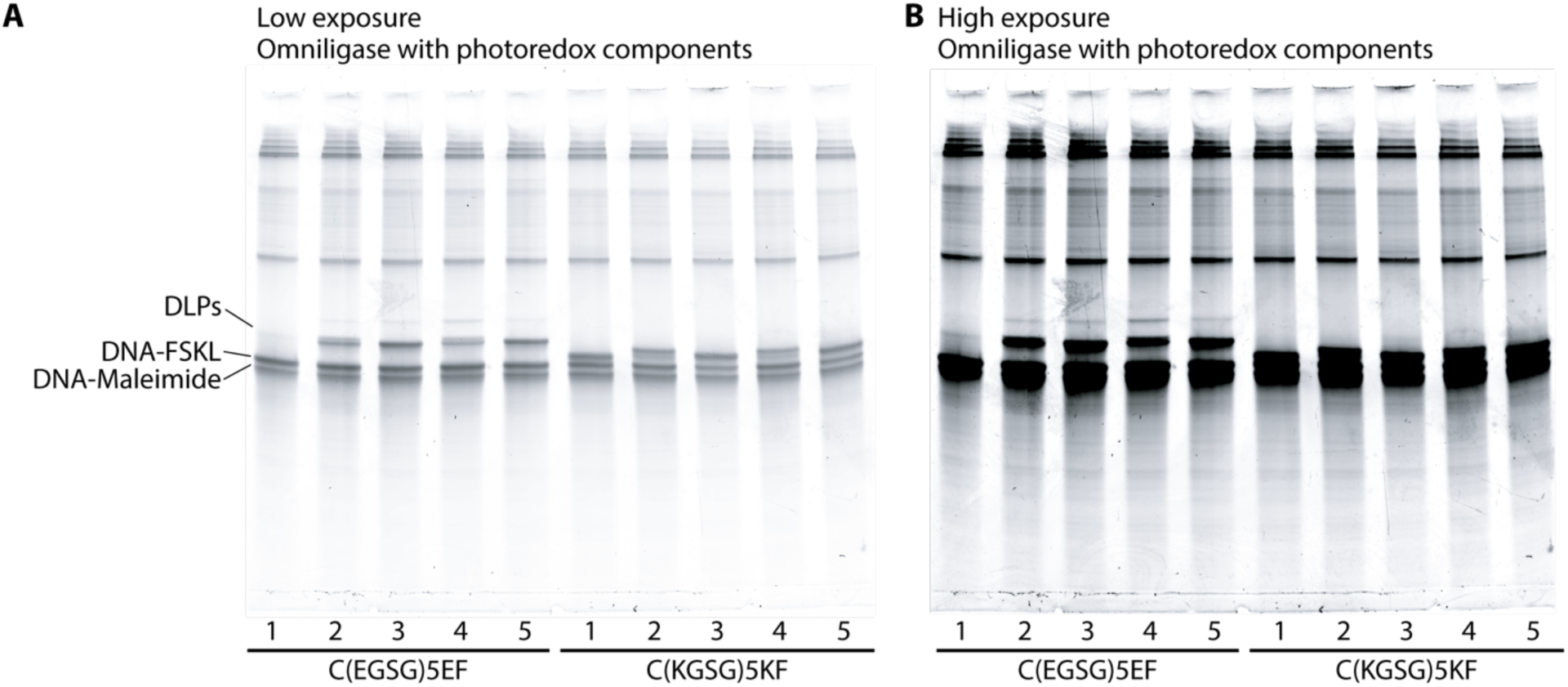
(A) Low-exposure and (B) high-exposure 10% Urea-PAGE gels comparing Omniligase ligation yields for peptides C(EGSG)₅EF and C(KGSG)₅KF after incubation with individual photoredox reaction components. Lanes: 1. 50 µM lumiflavin + 500 µM peptide in Milli-Q water, incubated for 3 h under 445 nm light (110 mW); 2. 2.5 mM NB–PEG₄–alkyne linker + 500 µM peptide in Milli-Q water, incubated for 3 h under 445 nm light (110 mW); 3. 10 mM Cs-formate (pH 3.5) + 500 µM peptide, incubated for 3 h under 445 nm light (110 mW); 4. 50 µM lumiflavin + 2.5 mM NB–PEG₄–alkyne linker + 10 mM Cs-formate (pH 3.5) + 500 µM peptide, incubated for 3 h at room temperature without irradiation; 5. 500 µM peptide in Milli-Q water, incubated for 3 h under 445 nm light (110 mW). All reactions for this gel were performed under ambient conditions (i.e. not in a nitrogen glove box). Gels were stained with SYBR Gold.

**Supplementary Figure 11.**
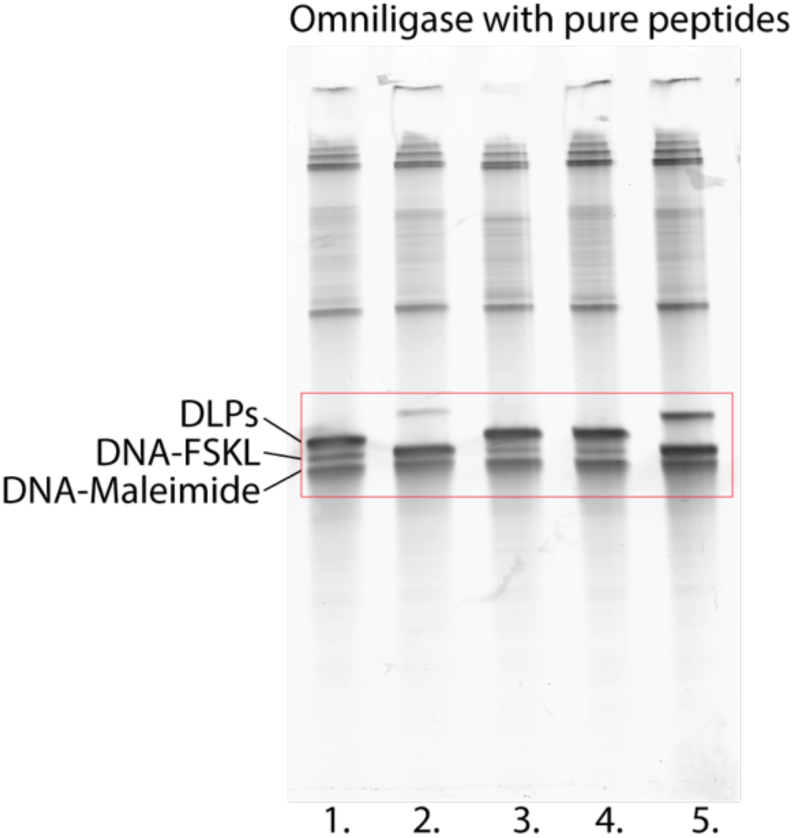
Uncropped 10% Urea-PAGE gel corresponding to Figure 2. The red rectangle indicates the region shown in the main figure. Lanes: 1. 1X FLAG; 2. 3X FLAG; 3. H3(1–21); 4. Ova(257–264); 5. β-Amyloid (1–16). Gel was stained with SYBR Gold.

**Supplementary Figure 12.**
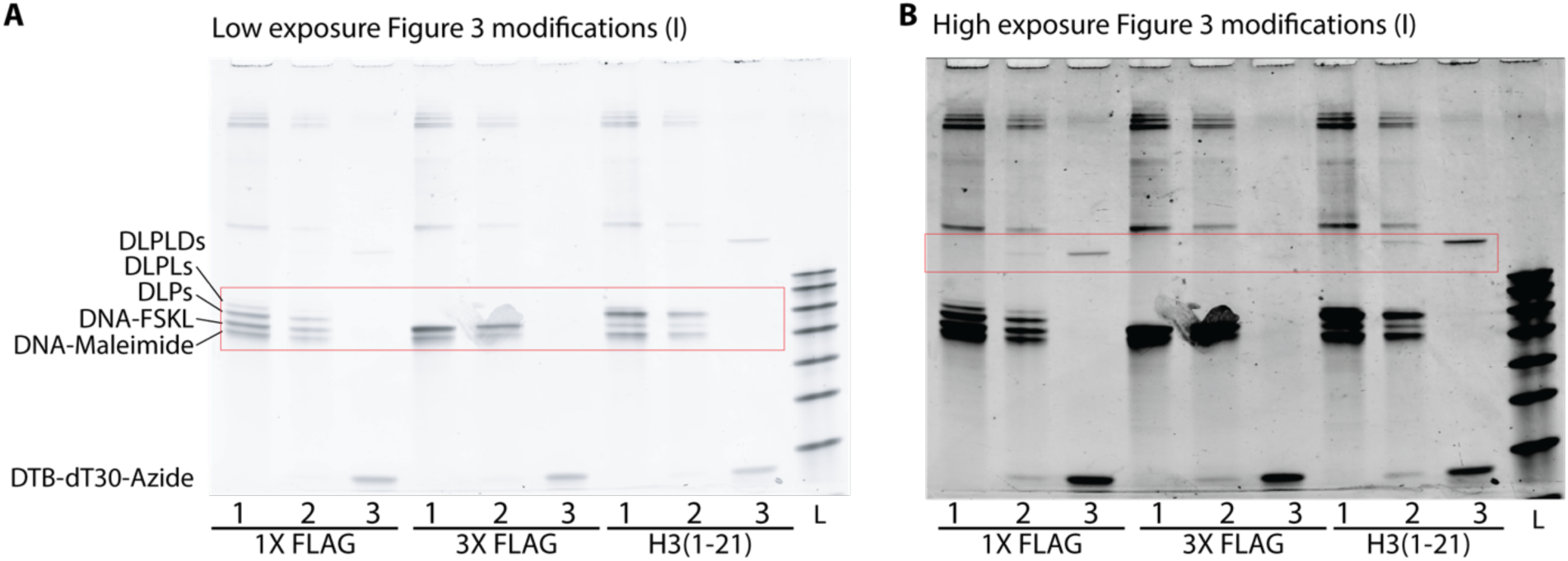
Uncropped 10% Urea-PAGE gels corresponding to Figure 3, shown at (A) low exposure and (B) high exposure. Lanes: 1. after photoredox modification and Omniligase ligation; 2. after CuAAC reaction; 3. after affinity purification with DynaBeads–Streptavidin to capture desthiobiotin-labeled dT₃₀ molecules; L. 20/100 ssDNA ladder. Red rectangles indicate regions shown in the main figure. Gel was stained with SYBR Gold.

**Supplementary Figure 13.**
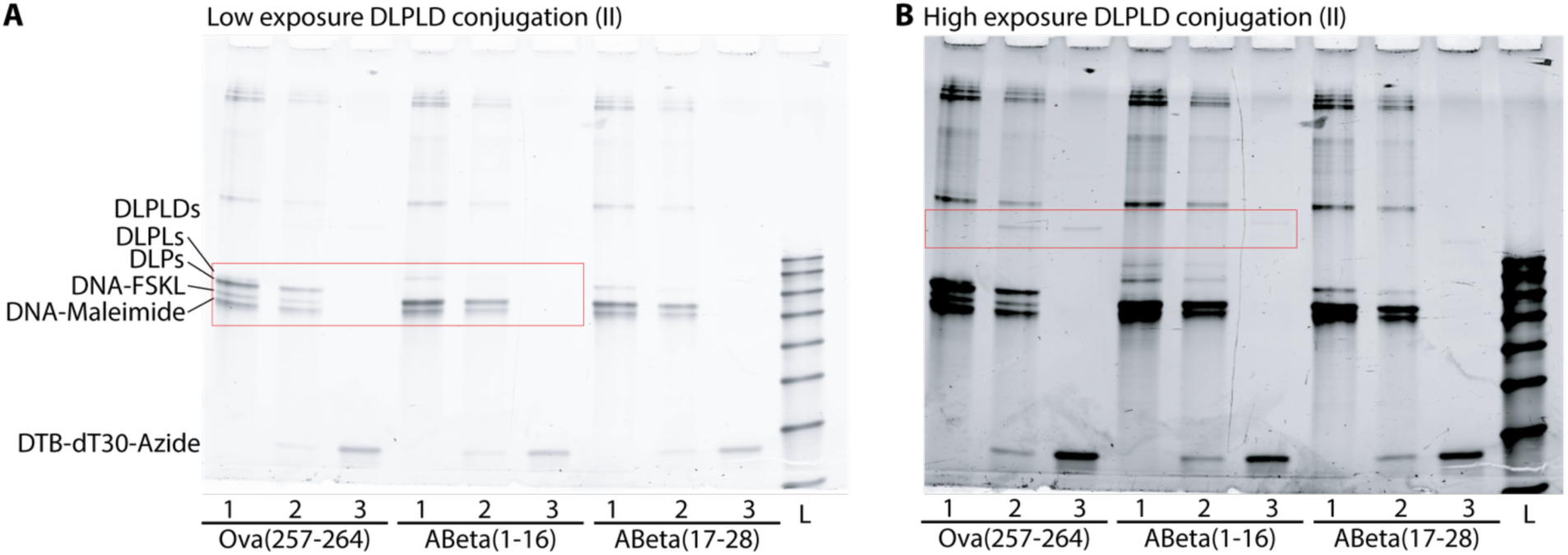
Uncropped 10% Urea-PAGE gels corresponding to Figure 3, shown at (A) low exposure and (B) high exposure. Lanes: 1. after photoredox modification and Omniligase ligation; 2. after CuAAC reaction; 3. after affinity purification with DynaBeads–Streptavidin to capture desthiobiotin-labeled dT₃₀ molecules; L. 20/100 ssDNA ladder. Red rectangles indicate regions shown in the main figure. Gels were stained with SYBR Gold.

**Supplementary Figure 14.**
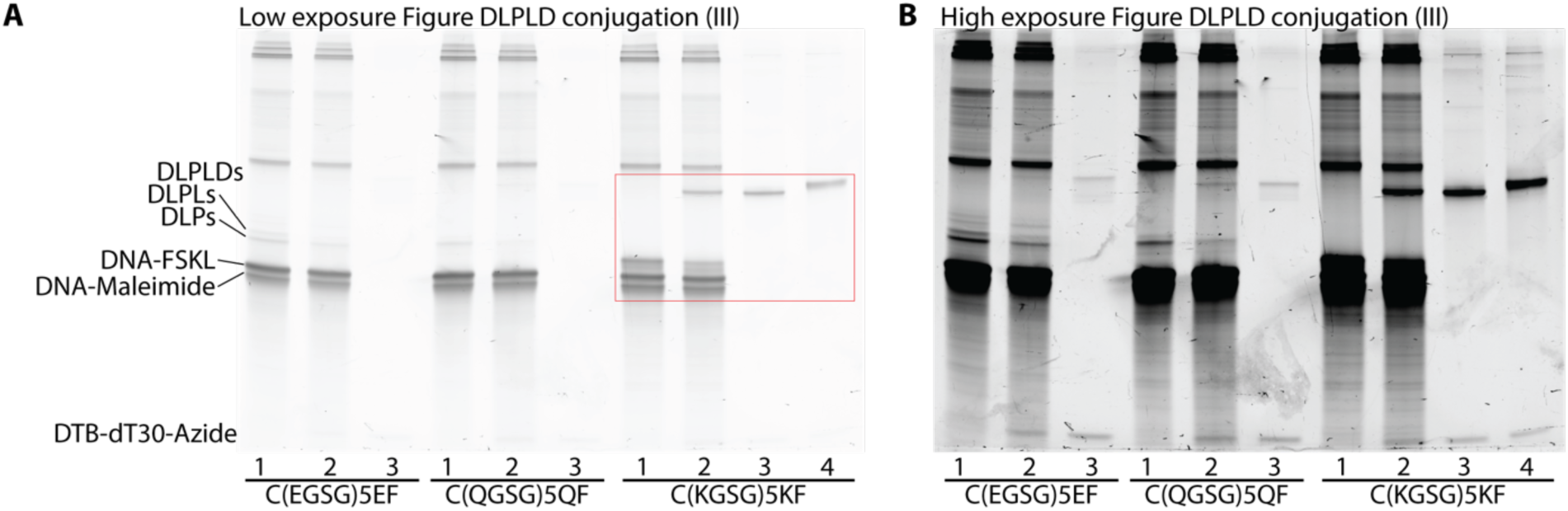
Uncropped 10% Urea-PAGE gels corresponding to Figure 3, shown at (A) low exposure and (B) high exposure. Lanes: 1. after photoredox modification and Omniligase ligation; 2. after CuAAC reaction; 3. after affinity purification with DynaBeads–Streptavidin to capture desthiobiotin-labeled dT₃₀ molecules; 4. after sulfo-NHS–acetylation; L. = 20/100 ssDNA ladder. Red rectangles indicate regions shown in the main figure. Gels were stained with SYBR Gold.

## References

(1) Deamer, D.; Akeson, M.; Branton, D. Three Decades of Nanopore Sequencing. Nat. Biotechnol. 2016, 34 (5), 518–524. 10.1038/nbt.3423.

(2) Mohammadi, M. M.; Bavi, O. DNA Sequencing: An Overview of Solid-State and Biological Nanopore-Based Methods. Biophys. Rev. 2021, 14 (1), 99–110. 10.1007/s12551-021-00857-y.

(3) Dorey, A.; Howorka, S. Nanopore DNA Sequencing Technologies and Their Applications towards Single-Molecule Proteomics. Nat. Chem. 2024, 16 (3), 314–334. 10.1038/s41557-023-01322-x.

(4) Alfaro, J. A.; Peggy R. Bohländer; Bohländer, P. R.; Dai, M.; Mike Filius; Filius, M.; Cecil J. Howard; Howard, C. J.; van Kooten, X. F.; Ohayon, S.; Adam Pomorski; Pomorski, A.; Schmid, S.; Schmid, S.; Sonja Schmid; Sonja Schmid; Aleksei Aksimentiev; Aksimentiev, A.; Anslyn, E. V.; Bedran, G.; Chan Cao; Chan Cao; Cao, C.; Chinappi, M.; Coyaud, E.; Dekker, C.; Dittmar, G.; Nicholas Drachman; Nicholas Drachman; Nicholas Drachman; Drachman, N.; Eelkema, R.; David R. Goodlett; Goodlett, D. R.; Hentz, S.; Kalathiya, U.; Kelleher, N. L.; Kelly, R. T.; Zvi Kelman; Kelman, Z.; Kim, D.; Sung Hyun Kim; Kim, S. H.; Rolain, J.-M.; Kuster, B.; Rodriguez-Larrea, D.; Lindsay, S.; Maglia, G.; Marcotte, E. M.; Marino, J. P.; Masselon, C.; Michael Mayer; Mayer, M.; Samaras, P.; Sarthak, K.; Sepiashvili, L.; Stein, D.; Wanunu, M.; Debrauwer, L.; Wilhelm, M.; Yin, P.; Meller, A.; Joo, C. The Emerging Landscape of Single-Molecule Protein Sequencing Technologies. Nat. Methods 2021, 18 (6), 604–617. 10.1038/s41592-021-01143-1.

(5) MacCoss, M. J.; Alfaro, J. A.; Faivre, D. A.; Wu, C. C.; Wanunu, M.; Slavov, N. Sampling the Proteome by Emerging Single-Molecule and Mass Spectrometry Methods. Nat. Methods 2023, 20 (3), 339–346. 10.1038/s41592-023-01802-5.

(6) Plesa, C.; Kowalczyk, S. W.; Zinsmeester, R.; Grosberg, A. Y.; Rabin, Y.; Dekker, C. Fast Translocation of Proteins through Solid State Nanopores. Nano Lett. 2013, 13 (2), 658–663. 10.1021/nl3042678.

(7) Manrao, E. A.; Derrington, I. M.; Laszlo, A. H.; Langford, K. W.; Hopper, M. K.; Gillgren, N.; Pavlenok, M.; Niederweis, M.; Gundlach, J. H. Reading DNA at Single-Nucleotide Resolution with a Mutant MspA Nanopore and Phi29 DNA Polymerase. Nat. Biotechnol. 2012, 30 (4), 349–353. 10.1038/nbt.2171.

(8) Cherf, G. M.; Lieberman, K. R.; Rashid, H.; Lam, C. E.; Karplus, K.; Akeson, M. Automated Forward and Reverse Ratcheting of DNA in a Nanopore at 5-Å Precision. Nat. Biotechnol. 2012, 30 (4), 344–348. 10.1038/nbt.2147.

(9) Nivala, J.; Marks, D. B.; Akeson, M. Unfoldase-Mediated Protein Translocation through an α-Hemolysin Nanopore. Nat. Biotechnol. 2013, 31 (3), 247–250. 10.1038/nbt.2503.

(10) Nivala, J.; Mulroney, L.; Li, G.; Schreiber, J.; Akeson, M. Discrimination among Protein Variants Using an Unfoldase-Coupled Nanopore. ACS Nano 2014, 8 (12), 12365–12375. 10.1021/nn5049987.

(11) Motone, K.; Kontogiorgos-Heintz, D.; Wee, J.; Kurihara, K.; Yang, S.; Roote, G.; Fox, O. E.; Fang, Y.; Queen, M.; Tolhurst, M.; Cardozo, N.; Jain, M.; Nivala, J. Multi-Pass, Single-Molecule Nanopore Reading of Long Protein Strands. Nature 2024, 1–8. 10.1038/s41586-024-07935-7.

(12) Cordova, J. C.; Olivares, A. O.; Shin, Y.; Stinson, B. M.; Calmat, S.; Schmitz, K. R.; Aubin-Tam, M.-E.; Baker, T. A.; Lang, M. J.; Sauer, R. T. Stochastic but Highly Coordinated Protein Unfolding and Translocation by the ClpXP Proteolytic Machine. Cell 2014, 158 (3), 647–658. 10.1016/j.cell.2014.05.043.

(13) Brinkerhoff, H.; Albert S W Kang; Jingqian Liu; Aksimentiev, A.; Dekker, C. Multiple Rereads of Single Proteins at Single-Amino Acid Resolution Using Nanopores. Science 2021. 10.1126/science.abl4381.

(14) Chen, Z.; Wang, Z.; Xu, Y.; Zhang, X.; Tian, B.; Bai, J. Controlled Movement of ssDNA Conjugated Peptide through Mycobacterium Smegmatis Porin A (MspA) Nanopore by a Helicase Motor for Peptide Sequencing Application. Chem. Sci. 2021, 12 (47), 15750–15756. 10.1039/D1SC04342K.

(15) Yan, S.; Zhang, J.; Wang, Y.; Guo, W.; Zhang, S.; Liu, Y.; Cao, J.; Wang, Y.; Wang, L.; Ma, F.; Zhang, P.; Chen, H.-Y.; Huang, S. Single Molecule Ratcheting Motion of Peptides in a Mycobacterium Smegmatis Porin A (MspA) Nanopore. Nano Lett. 2021, 21 (15), 6703–6710. 10.1021/acs.nanolett.1c02371.

(16) Craig, J. M.; Laszlo, A. H.; Brinkerhoff, H.; Derrington, I. M.; Noakes, M. T.; Nova, I. C.; Tickman, B. I.; Doering, K.; de Leeuw, N. F.; Gundlach, J. H. Revealing Dynamics of Helicase Translocation on Single-Stranded DNA Using High-Resolution Nanopore Tweezers. Proc. Natl. Acad. Sci. 2017, 114 (45), 11932–11937. 10.1073/pnas.1711282114.

(17) Ian C. Nova; Justas Ritmejeris; H. Brinkerhoff; Theo J R Koenig; J. Gundlach; C. Dekker. Detection of Phosphorylation Post-Translational Modifications along Single Peptides with Nanopores. Nat. Biotechnol. 2023. 10.1038/s41587-023-01839-z.

(18) Chen, X.; van de Sande, J. W.; Ritmejeris, J.; Wen, C.; Brinkerhoff, H.; Laszlo, A. H.; Albada, B.; Dekker, C. Resolving Sulfation Posttranslational Modifications on a Peptide Hormone Using Nanopores. ACS Nano 2024, 18 (42), 28999–29007. 10.1021/acsnano.4c09872.

(19) Chen, Z.; Wang, J.; Meng, X.; Ye, F.; Wang, L.; Xu, Y.; Wang, X.; Wang, Q.; Wen, H.; Bai, J. Spike Signals and MD Simulations Reveal the Significance of Peptide Stretching in Nanopore Protein Sequencing. J. Am. Chem. Soc. 2025. 10.1021/jacs.5c00827.

(20) De Rosa, L.; Di Stasi, R.; Romanelli, A.; D’Andrea, L. D. Exploiting Protein N-Terminus for Site-Specific Bioconjugation. Molecules 2021, 26 (12), 3521. 10.3390/molecules26123521.

(21) Dawson, E. P.; Muir, T. W.; Ian, C.-L.; Kent B. H. Stephen. Synthesis of Proteins by Native Chemical Ligation. 10.1126/science.7973629.

(22) Thapa, P.; Zhang, R.-Y.; Menon, V.; Bingham, J.-P. Native Chemical Ligation: A Boon to Peptide Chemistry. Molecules 2014, 19 (9), 14461–14483. 10.3390/molecules190914461.

(23) Scheck, R. A.; Dedeo, M. T.; Iavarone, A. T.; Francis, M. B. Optimization of a Biomimetic Transamination Reaction. J. Am. Chem. Soc. 2008, 130 (35), 11762–11770. 10.1021/ja802495w.

(24) Dempsey, D. R.; Jiang, H.; Kalin, J. H.; Chen, Z.; Cole, P. A. Site-Specific Protein Labeling with N-Hydroxysuccinimide-Esters and the Analysis of Ubiquitin Ligase Mechanisms. J. Am. Chem. Soc. 2018, 140 (30), 9374–9378. 10.1021/jacs.8b05098.

(25) MacDonald, J. I.; Munch, H. K.; Moore, T.; Francis, M. B. One-Step Site-Specific Modification of Native Proteins with 2-Pyridinecarboxyaldehydes. Nat. Chem. Biol. 2015, 11 (5), 326–331. 10.1038/nchembio.1792.

(26) Filius, M.; van Wee, R.; de Lannoy, C.; Westerlaken, I.; Li, Z.; Kim, S. H.; de Agrela Pinto, C.; Wu, Y.; Boons, G.-J.; Pabst, M.; de Ridder, D.; Joo, C. Full-Length Single-Molecule Protein Fingerprinting. Nat. Nanotechnol. 2024, 1–8. 10.1038/s41565-023-01598-7.

(27) Bridge, H. N.; Leiter, W.; Frazier, C. L.; Weeks, A. M. An N Terminomics Toolbox Combining 2-Pyridinecarboxaldehyde Probes and Click Chemistry for Profiling Protease Specificity.

(28) Barber, L. J.; Stankevich, K. S.; Spicer, C. D. Effect of Pyridinecarboxaldehyde Functionalization on Reactivity and N-Terminal Protein Modification. ACS Publ. 2025. 10.1021/jacsau.5c00238.

(29) Guimaraes, C. P.; Witte, M. D.; Theile, C. S.; Bozkurt, G.; Kundrat, L.; Blom, A. E. M.; Ploegh, H. L. Site-Specific C-Terminal Internal Loop Labeling of Proteins Using Sortase-Mediated Reactions. Nat. Protoc. 2013, 8 (9), 1787–1799. 10.1038/nprot.2013.101.

(30) Antos, J. M.; Truttmann, M. C.; Ploegh, H. L. Recent Advances in Sortase-Catalyzed Ligation Methodology. Curr. Opin. Struct. Biol. 2016, 38, 111–118. 10.1016/j.sbi.2016.05.021.

31. Niemeyer, C. M. Semisynthetic DNA–Protein Conjugates for Biosensing and Nanofabrication. Angew. Chem. Int. Ed. 2010, 49 (7), 1200–1216. 10.1002/anie.200904930.

32. Williams, B. A. R.; Chaput, J. C. Synthesis of Peptide-Oligonucleotide Conjugates Using a Heterobifunctional Crosslinker. Curr. Protoc. Nucleic Acid Chem. Ed. Serge Beaucage Al 2010, CHAPTER, Unit4.41. 10.1002/0471142700.nc0441s42.

(33) Schmidt, M.; Toplak, A.; Quaedflieg, P. J. L. M.; Ippel, H.; Richelle, G. J. J.; Hackeng, T. M.; van Maarseveen, J. H.; Nuijens, T. Omniligase-1: A Powerful Tool for Peptide Head-to-Tail Cyclization. Adv. Synth. Catal. 2017, 359 (12), 2050–2055. 10.1002/adsc.201700314.

(34) Toplak, A.; Teixeira de Oliveira, E. F.; Schmidt, M.; Rozeboom, H. J.; Wijma, H. J.; Meekels, L. K. M.; de Visser, R.; Janssen, D. B.; Nuijens, T. From Thiol-Subtilisin to Omniligase: Design and Structure of a Broadly Applicable Peptide Ligase. Comput. Struct. Biotechnol. J. 2021, 19, 1277–1287. 10.1016/j.csbj.2021.02.002.

(35) Bloom, S.; Liu, C.; Kölmel, D. K.; Qiao, J. X.; Zhang, Y.; Poss, M. A.; Ewing, W. R.; MacMillan, D. W. C. Decarboxylative Alkylation for Site-Selective Bioconjugation of Native Proteins via Oxidation Potentials. Nat Chem 2018, 10 (2), 205–211. 10.1038/nchem.2888.

(36) Zhang, L.; Floyd, B. M.; Chilamari, M.; Mapes, J.; Swaminathan, J.; Bloom, S.; Marcotte, E. M.; Anslyn, E. V. Photoredox-Catalyzed Decarboxylative C-Terminal Differentiation for Bulk- and Single-Molecule Proteomics. ACS Chem. Biol. 2021, 16 (11), 2595–2603. 10.1021/acschembio.1c00631.

37. V, H.; Nf, S.; M, M.; Mg, F. Labeling Live Cells by Copper-Catalyzed Alkyne--Azide Click Chemistry. PubMed 2010.

(38) Zapadka, K. L.; Becher, F. J.; Gomes dos Santos, A. L.; Jackson, S. E. Factors Affecting the Physical Stability (Aggregation) of Peptide Therapeutics. Interface Focus 2017, 7 (6), 20170030. 10.1098/rsfs.2017.0030.

(39) Li, R.; Schmidt, M.; Zhu, T.; Yang, X.; Feng, J.; Tian, Y.; Cui, Y.; Nuijens, T.; Wu, B. Traceless Enzymatic Protein Synthesis without Ligation Sites Constraint. Natl. Sci. Rev. 2022, 9 (5), nwab158. 10.1093/nsr/nwab158.

(40) Gimeno, A.; Ehlers, A. M.; Delgado, S.; Langenbach, J.-W. H.; van den Bos, L. J.; Kruijtzer, J. A. W.; Guigas, B. G. A.; Boons, G.-J. Site-Specific Glyco-Tagging of Native Proteins for the Development of Biologicals. J. Am. Chem. Soc. 2024, 146 (50), 34452–34465. 10.1021/jacs.4c11091.

(41) Lu, C.; Bonini, A.; Viel, J. H.; Maglia, G. Toward Single-Molecule Protein Sequencing Using Nanopores. Nat. Biotechnol. 2025, 43 (3), 312–322. 10.1038/s41587-025-02587-y.

(42) Sauciuc, A.; Whittaker, J.; Tadema, M.; Tych, K.; Guskov, A.; Maglia, G. Blobs Form during the Single-File Transport of Proteins across Nanopores. Proc. Natl. Acad. Sci. 2024, 121 (38), e2405018121. 10.1073/pnas.2405018121.

43. François Petitjean; Alain Ketterlin; Pierre Gançarski. A Global Averaging Method for Dynamic Time Warping, with Applications to Clustering. 2011. 10.1016/j.patcog.2010.09.013.

(44) Asandei, A.; Schiopu, I.; Chinappi, M.; Seo, C. H.; Park, Y.; Luchian, T. Electroosmotic Trap Against the Electrophoretic Force Near a Protein Nanopore Reveals Peptide Dynamics During Capture and Translocation. ACS Appl. Mater. Interfaces 2016, 8 (20), 13166–13179. 10.1021/acsami.6b03697.

(45) Huang, G.; Willems, K.; Soskine, M.; Wloka, C.; Maglia, G. Electro-Osmotic Capture and Ionic Discrimination of Peptide and Protein Biomarkers with FraC Nanopores. Nat. Commun. 2017, 8 (1), 935. 10.1038/s41467-017-01006-4.

(46) Restrepo-Pérez, L.; Wong, C. H.; Maglia, G.; Dekker, C.; Joo, C. Label-Free Detection of Post-Translational Modifications with a Nanopore. Nano Lett. 2019, 19 (11), 7957–7964. 10.1021/acs.nanolett.9b03134.

(47) Yu, L.; Kang, X.; Li, F.; Mehrafrooz, B.; Makhamreh, A.; Fallahi, A.; Foster, J. C.; Aksimentiev, A.; Chen, M.; Wanunu, M. Unidirectional Single-File Transport of Full-Length Proteins through a Nanopore. Nat. Biotechnol. 2023, 41 (8), 1130–1139. 10.1038/s41587-022-01598-3.

(48) Sauciuc, A.; Morozzo della Rocca, B.; Tadema, M. J.; Chinappi, M.; Maglia, G. Translocation of Linearized Full-Length Proteins through an Engineered Nanopore under Opposing Electrophoretic Force. Nat. Biotechnol. 2023, 1–7. 10.1038/s41587-023-01954-x.

49. P Martin-Baniandres; WH Lan; S Board; M Romero-Ruiz; S Garcia-Manyes; Y Qing; H Bayley. Enzyme-Less Nanopore Detection of Post-Translational Modifications within Long Polypeptides. Nat. Nanotechnol. 2023. 10.1038/s41565-023-01462-8.

## Supplementary references

(1) Chen, X.; van de Sande, J. W.; Ritmejeris, J.; Wen, C.; Brinkerhoff, H.; Laszlo, A. H.; Albada, B.; Dekker, C. Resolving Sulfation Posttranslational Modifications on a Peptide Hormone Using Nanopores. ACS Nano 2024, 18 (42), 28999–29007. 10.1021/acsnano.4c09872.

(2) Zhang, L.; Floyd, B. M.; Chilamari, M.; Mapes, J.; Swaminathan, J.; Bloom, S.; Marcotte, E. M.; Anslyn, E. V. Photoredox-Catalyzed Decarboxylative C-Terminal Differentiation for Bulk- and Single-Molecule Proteomics. ACS Chem Biol 2021, 16 (11), 2595–2603. 10.1021/acschembio.1c00631.

(3) Aquilina, M.; Wu, N. J. W.; Kwan, K.; Bušić, F.; Dodd, J.; Nicolás-Sáenz, L.; O’Callaghan, A.; Bankhead, P.; Dunn, K. E. GelGenie: An AI-Powered Framework for Gel Electrophoresis Image Analysis. Nature Communications 2025, 16 (1), 4087. 10.1038/s41467-025-59189-0.

